# A Dual-Ensemble Model of Infralimbic Cortex Function in Behavioural Flexibility

**DOI:** 10.64898/2026.06.29.735227

**Authors:** Avi Adlakha, Ivo Sonntag, Simone Pfarr, Janet Barroso-Flores, Wolfgang Sommer, Thomas Kuner

**Author notes:** Equal contribution (Wolfgang Sommer, Thomas Kuner).

## Abstract

The infralimbic cortex (IL) has been described as both facilitator and terminator of reward-seeking, yet both functions appear mutually exclusive. We reconcile this discrepancy by proposing a dual-ensemble model: one for action-outcome prediction and another for prediction correction. Using longitudinal miniscope imaging in rats, we show that during training, the action-outcome predictor ensemble sharpens its activity by suppressing the majority of the IL neuronal population. As contingencies stabilize, entropy decreases, reflecting a refined predictive model. During extinction, the prediction correction ensemble activates in response to reward omission. A computational simulation based on these two ensembles successfully recapitulated our experimental results. This framework reconciles conflicting models, where IL manipulation can either accelerate or abolish extinction by demonstrating that IL activity dynamically encodes predictive accuracy. Our findings indicate that IL functions as an action-outcome predictor that elegantly guides behavioural flexibility.

## Introduction

Adaptive behaviour depends on the ability to predict the consequences of actions and to update these predictions when environmental contingencies change. The medial prefrontal cortex (mPFC) plays a central role in this process, integrating motivational, contextual, and action-related information to guide flexible decision-making ^1^. The functional landscape of the infralimbic (IL) cortex has historically been defined by its role in fear extinction, particularly through foundational studies from the Quirk laboratory ^2–10^. These early ablation and lesion studies established the IL as an essential “STOP” region for behaviour, demonstrating that while the IL is not required to learn a fear task, it is critical for suppressing freezing behaviour. Consequently, the region was long viewed as a dedicated inhibitory switch. However, this model failed to explain findings in drug and food seeking behaviours. While some findings supported an extinction role, others revealed that IL inhibition slowed the acquisition of self-administration, suggesting a facilitatory function ^11–17^. Theories attempting to bridge this gap, formulating context differences or movement-based similarities, could not reconcile this discrepancy ^18–22^. The IL cortex is also deeply involved in habit formation, specifically the transition from goal directed actions to stimulus response patterns ^23–25^. This introduces a functional paradox; if the IL is necessary for habit formation, its removal should theoretically accelerate extinction by preventing habit repetition, yet the opposite occurs.

A key challenge in interpreting these findings is that the functional contribution of the IL appears highly context dependent. Experimental manipulations of IL activity have yielded opposing behavioural outcomes, with disruption either impairing extinction or enhancing reward seeking depending on task and timing ^14,26–28^, suggesting that IL function cannot be adequately described as a unitary process. Instead, it raises the possibility that distinct neuronal subpopulations within the IL support different, potentially opposing computations, whose relative engagement depends on behavioural demands ^22^. Recent studies have begun to identify specific ensembles corresponding to these distinct functions, demonstrating that targeted manipulations of these assemblies can selectively impair one behavioural facet while leaving others intact ^17,29^.

From a computational perspective, this apparent functional duality is reminiscent of frameworks in which cortical circuits encode both predictions about action outcomes and deviations from those predictions. In particular, models of medial prefrontal function propose that neural populations represent expected outcomes while parallel populations signal prediction corrections that drive behavioural updating ^30,31^. Such architectures provide a mechanism for balancing behavioural persistence with flexibility, yet their relevance to IL function during appetitive learning has not been directly tested.

Here, we propose that the IL implements a dual-ensemble architecture comprising (i) an *action–outcome predictor ensemble,* which encodes expected consequences of behaviour, and (ii) a *prediction correction ensemble*, which is recruited when outcomes deviate from expectation. Under this framework, several key observations arise as necessary consequences of its operation including the sparse population of responding neurons ^26,29^ and decreasing neural recruitment by repeated actions and habit formation ^23,25,32^. Most importantly, the high activity levels and necessary contribution of the IL cortex during extinction ^6,33^ are revealed to be an inherent by-product of the framework’s underlying mechanisms. This hypothesis offers a unifying account that reconciles previously conflicting findings by linking IL activity to predictive dual-ensemble model.

To test this, we performed longitudinal calcium imaging of neuronal populations in the IL cortex of freely behaving rats using head-mounted miniscopes during a cue-conditioned operant task encompassing acquisition, extinction and reinstatement. This approach enables tracking of single neurons and population dynamics across task conditions with high temporal resolution ^34^. We find that IL activity is organized into dynamically evolving ensembles that are not rigidly tied to specific behavioural events but instead exhibit flexible temporal tuning across trials. As learning progresses, population activity becomes increasingly structured, reflected in a robust decrease in Shannon’s entropy both within and across sessions. Entropy reduction indicates a transition from exploratory, high-variability dynamics to more stable, predictive representations. Concomitantly, ensemble structure reveals stable core populations embedded within more dynamic neuronal assemblies, consistent with the emergence of refined internal models of task contingencies.

## Results

### Rats learn and adapt reward-seeking strategies across self-administration, extinction, and reinstatement

To examine how behavioural strategies and outcome expectations evolve across self-administration, extinction, and reinstatement, rats were equipped with head-mounted miniscopes to record neuronal activity in the infralimbic cortex (IL) and subsequently trained in a cue-conditioned self-administration task (**Fig. 1A; Supplementary Fig. 1A-C**). Recordings throughout the self-administration phase could be obtained from five rats, of which three also completed the extinction and reinstatement task conditions. During self-administration, rats rapidly acquired the task, exhibiting a strong preference for the active lever (68.9± 8.4 presses/session) over the inactive lever (2.2 ± 0.5 presses/session, p=6.6e-10), indicating robust action–outcome learning (**Fig. 1B; Supplementary Fig. 1D**). Lever pressing initially increased across sessions and subsequently declined, consistent with the incorporation of task structure, including reward delivery and timeout intervals. Following removal of both cue and reward, active lever pressing decreased markedly during extinction (8.4 ± 2.0 presses/session; p=0.016), indicating suppression of the learned behaviour. In the reinstatement task condition, reintroduction of the cue without reward led to a significant recovery of responding (13.7 ± 2.7 presses/session; p=0.005), although behaviour remained below self-administration levels (**Fig. 1B; Supplementary Fig. 1D**).

**Figure 1:**
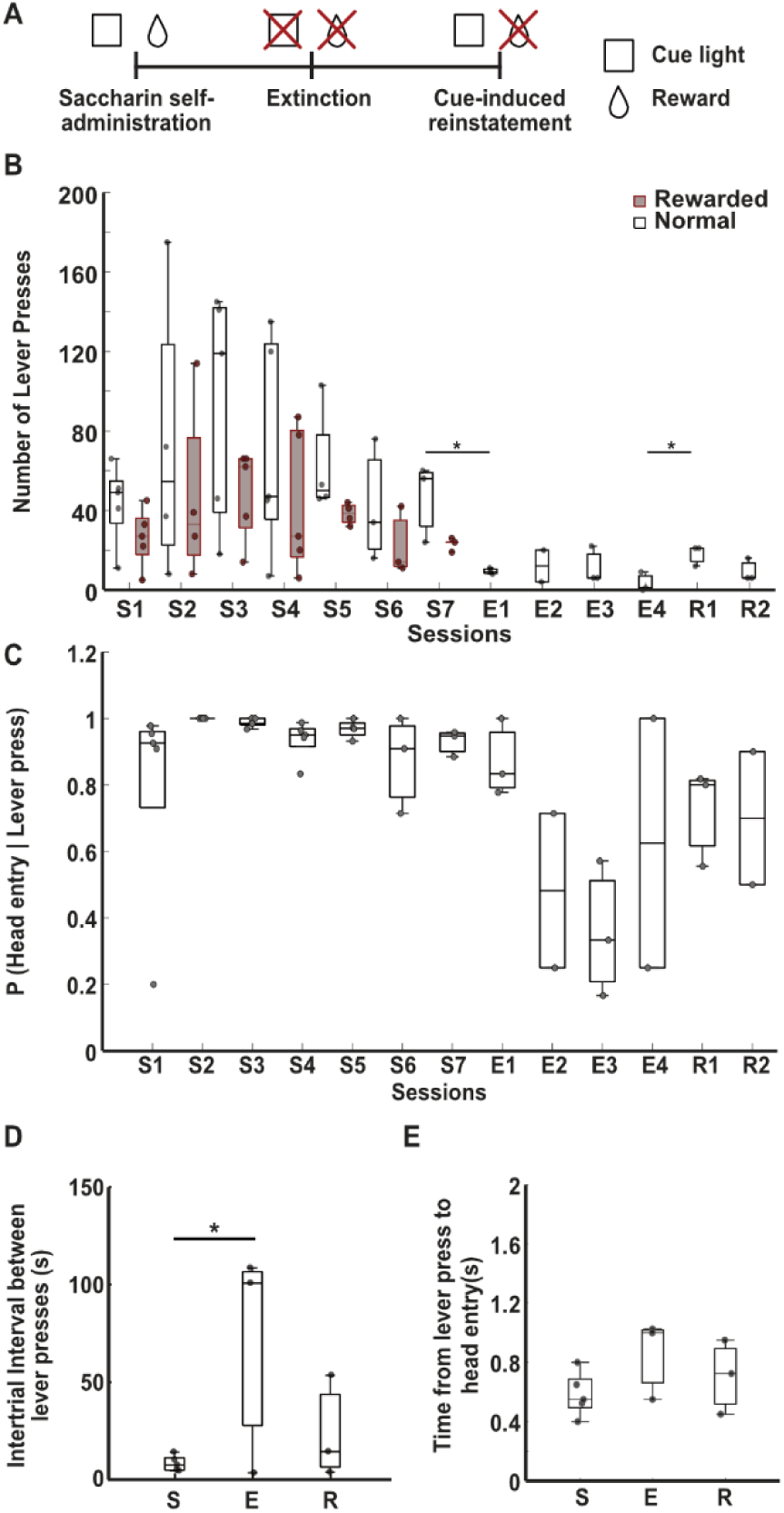
Behavioural characterization of cue-conditioned operant task. **A.** Schematic of experimental design showing self-administration (S, reward + cue), extinction (E, no reward, no cue), and reinstatement (R, cue only). **B.** Lever presses (total: black; rewarded: red) across sessions. **C.** Probability of a head entry following a lever press. **D.** Inter-trial interval (ITI) between consecutive active lever presses across task conditions. **E.** Latency from lever press to subsequent head entry. For the box plots, the central mark indicates the median, and the bottom and top edges of the box indicate the 25th and 75th percentiles, respectively. The whiskers extend to the most extreme data points not considered outliers. Each individual scatter point represents an animal. (for S n=5, for E,R n=3), *p<0.05 Liner-mixed effects model.

To quantify reward expectation independently of lever pressing, we analysed head-entry behaviour following trial initiation. During self-administration, rats performed a head entry following 93% of lever presses, reflecting a high expectation of reward availability (**Fig. 1C; Supplementary Fig. 1E**). This probability decreased to 57% during extinction (p=3.5e-05 vs Self-administration), indicating reduced expectation in the absence of predictive cues. Notably, reinstatement restored this measure to 76.2% (p=0.049 vs Extinction), demonstrating that the cue alone is sufficient to (partially) reinstate reward expectation despite continued reward omission. Consistent with this interpretation, the inter-trial interval (ITI) increased substantially during extinction (49.6 ± 19.8 s vs 16.2 ± 5.5 s during self-administration; p= 0.0058), reflecting reduced motivation for the task. During reinstatement, ITIs decreased to 23.9 ± 15.8 s, indicating partial recovery of task engagement (**Fig. 1D; Supplementary Fig. 1F**), although this effect did not reach significance relative to other task conditions (p = 0.06 vs extinction, and 0.72 vs self-administration).

In contrast, the latency between lever press and head entry remained stable across all task conditions (∼700ms; p = 0.82; **Fig. 1E; Supplementary Fig. 1G**), suggesting that the motor sequence linking action and outcome evaluation is highly automated and insensitive to changes in reward contingency. However, the number and cumulative duration of head entries per session was significantly modulated by task condition (56.6 ± 10.9 s, 133.52 ± 20.6 s during self-administration, 4 ± 0.4 s, 3.45 ± 1.6 s during extinction, and 11.8 ± 3.4 s, 16.2 ± 5.9 s during reinstatement, with p = 2.38e-07 and 1.90e-19 respectively), reflecting adaptive adjustment to expected reward availability (**Supplementary Fig.1H-K**).

Together, these results demonstrate that the behavioural paradigm induces robust learning and adaptive modification of reward-seeking strategies. Cue-induced reinstatement restores reward expectation and task engagement, as reflected by increased outcome checking and reduced inter-trial intervals. The distinct behavioural features provide a framework for probing how IL neuronal ensembles encode distinct components of action–outcome representations and their dynamic updating across task conditions.

### In-vivo calcium imaging reveals behaviourally aligned IL neuronal activity

To illustrate the quality and structure of the recorded neuronal activity, we present representative calcium imaging data from a single animal’s first session (Rat 1, Self-Admin. 01). Maximum intensity projections (MIPs) of raw and processed recordings demonstrate the extraction of neuronal signals and the identification of individual somata following pre-processing **(Fig. 2 A, B)**. Spatial footprints of identified neurons are shown in **Fig. 2C**, and histological verification confirmed accurate GRIN lens placement and viral expression within the infralimbic cortex **(Fig. 2D)**. The spatial (Fig. 2C) and temporal (Fig. 2E) footprints of five randomly selected neurons are highlighted in blue and with arrows to illustrate representative activity. The analysis pipeline yielded high-quality signals with a high signal-to-noise ratio, demonstrating effective separation of neuronal activity from background and motion artifacts. These calcium transients exhibited characteristic kinetics, defined by a rapid onset followed by a slow decay phase lasting several seconds and pronounced event-related transients, illustrating the diversity of response profiles within the recorded population (**Fig. 2F**). Importantly, these examples are representative of the broader dataset and serve to qualitatively demonstrate the presence of behaviourally aligned neuronal activity in IL ensembles. A systematic and quantitative analysis of trial-related tuning and population dynamics across animals and sessions is presented in the following section.

**Figure 2.**
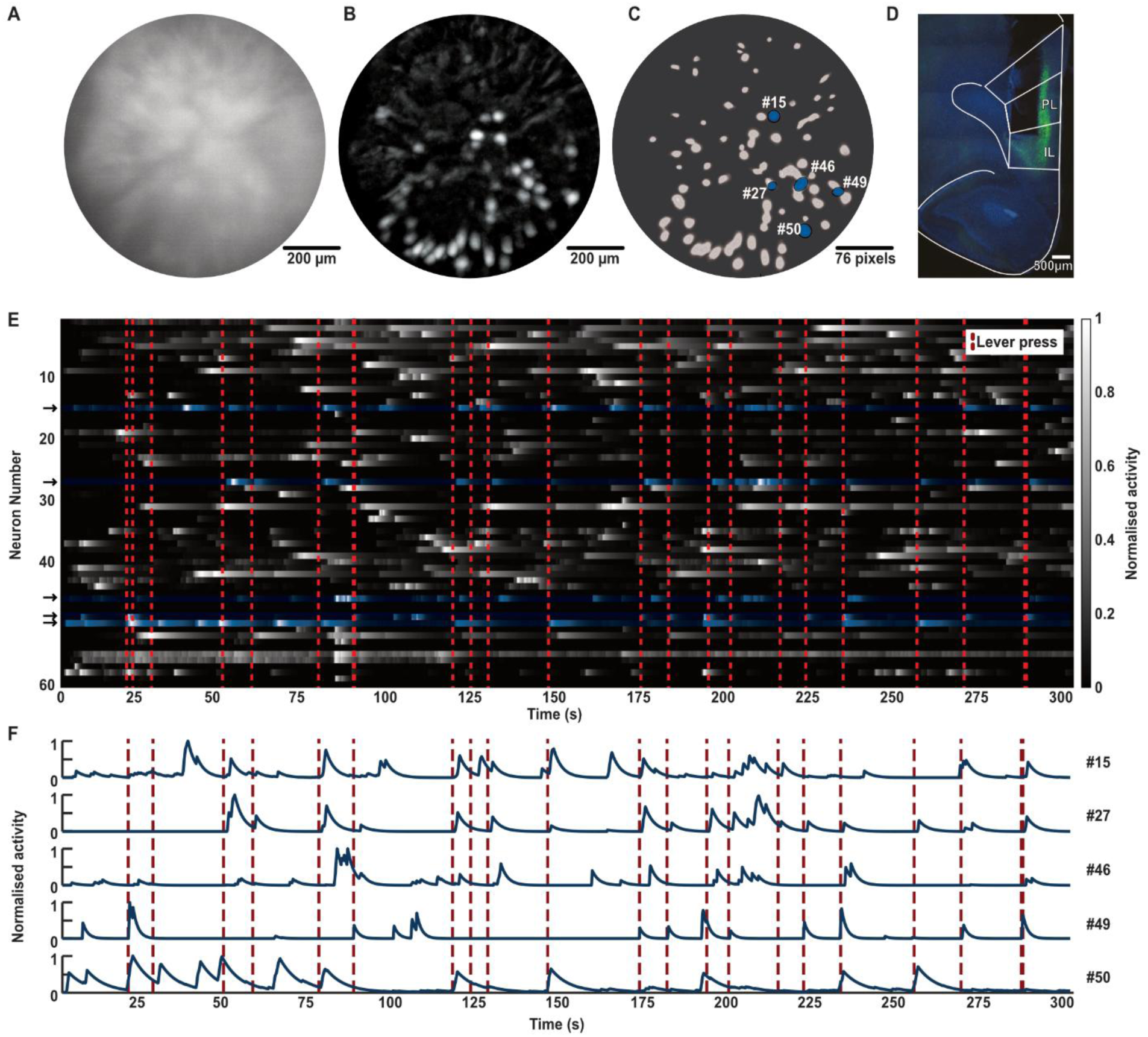
In vivo calcium imaging and extraction of neural dynamics in the infralimbic cortex. **A.** Maximum intensity projection (MIP) of raw fluorescence during the initial 5 minutes of a representative self-administration session (Rat 1, S1); scale bar, 200um. **B.** Background-subtracted (dF/F) MIP for the same epoch, illustrating enhanced signal-to-noise contrast; scale bar, 200um. **C.** Extracted spatial footprints of identified neurons within the field of view (FOV). Five representative neurons are highlighted in blue for subsequent representation; scale bar, 76 pixels. **D.** Histological verification of GRIN lens placement targeting the infralimbic (IL) cortex. Immunofluorescence showing GCaMP6 expression (green) and DAPI nuclear staining (blue). **E.** Temporal activity heatmap for 60 neurons over the 5-minute period (Rat 1, SA1). Neuronal activity is normalized (0–1, grayscale) and plotted against time. Red dashed lines denote lever presses. Five representative neurons are highlighted in blue. **F.** Extracted calcium traces for the five representative neurons highlighted in (C) and (E), showing normalized fluorescence intensity across the 5-minute representative recording window.

### Sparse and temporally structured IL activity is dominated by trial-related suppression

To characterize how IL neurons encode trial-related events, we quantified calcium activity in a peri-event window spanning −5 s to +5 s around the lever press. Neuronal activity was compared against a bootstrapped baseline to determine significant modulation (See Methods). Notably, some IL neurons exhibited highly tuned responses, with their activity largely confined to specific peri-event periods. A representative example is shown in **Fig. 3A**, where trial-by-trial heatmaps (**Fig. 3A1**) and the corresponding mean response (**Fig. 3A2**) reveal a reliable, temporally confined activation pattern aligned to the behavioural sequence. These tuned responses were individualistic and confined to specific parts of the task for each neuron during the acquisition of the task. **Figure 3B** further illustrates this with three distinct examples: one with peak activity preceding the lever press, another peaking after the lever press but before head entry, and a third activating after head entry. The neurons activity modulation was not just limited to being excited, but they also showed suppression correlating with the lever press. An example of a neuron exhibiting significant suppression is shown in **Fig. 3C**. The percentage of suppressed neurons during self-administration and extinction (41.0%, 34.2%) was found to be significantly higher than that of activated neurons (7.4%, 6.8%; p= 0.0001 for both). However, the percentage of activated neurons went up significantly during the reinstatement sessions (21.3%, p=0.006), although the percentage of suppressed neurons remained stable (32.4%, p=0.81) (**Fig. 3D**) The increased percentage of activated reinstatement neurons, motivated us to look into the ensembles that appear during the different session types, and their activity signatures. This imbalance indicates that trial-related suppression is a stable and prominent feature of IL population dynamics with a higher recruitment of activated neurons during reinstatement.

**Figure 3:**
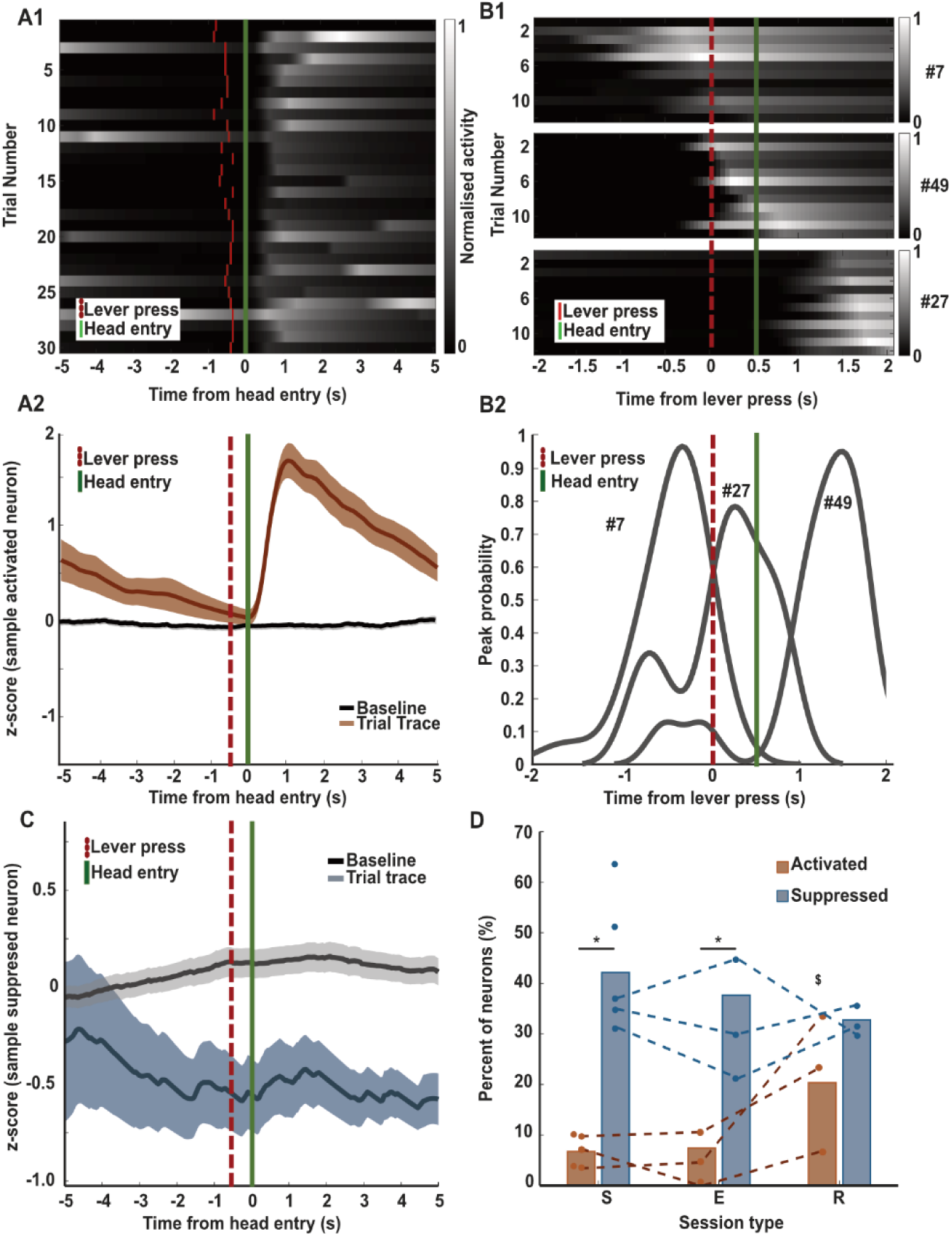
Tuned Cells and their activity around behavioural markers. **A1.** Representative heatmap of an activated neuron time-locked to the head entry (green line). Each row represents a single trial; the red vertical line denotes the preceding lever press. **A2.** A line graph showing the mean z-scored activity for all trials (orange dark line) for a sample activated neuron. Shaded area represents ± SEM. The grey horizontal line indicates the threshold for significance derived from bootstrapped null distributions. **B1**. Trial-by-trial heatmaps for three functional cell types: neurons showing peak activity prior to a lever press (pre-LP), between the lever press and head entry (LP-HE interval), and following a head entry (post-HE). **B2.** Probability density distributions of peak activity relative to behavioural markers for the cell types shown in (D1), calculated across all recorded trials. **C.** Line graph representative suppressed neuron showing mean z-scored activity (blue dark line) relative to the behavioural event; shaded area represents ± SEM. **D.** Percentage of significantly activated (orange) and suppressed (blue) neurons across self-administration, extinction, and reinstatement sessions. * p-value<0.05, $ activated neuron percentage in R sig. different from SA and E sessions, p= 0.0097.

Together, these findings indicate that IL ensemble activity during goal-directed behaviour is characterized by sparse and temporally structured excitation embedded within widespread suppression. This pattern suggests that task-relevant information is encoded by a small subset of selectively active neurons against a background of reduced population activity, consistent with a high signal-to-noise coding regime. Such an organization is compatible with computational frameworks in which predictive representations are sharpened through selective activation and concurrent suppression of non-informative activity ^35,36^.

### Structured IL population dynamics differentiate behavioural contingencies

Analysis of z-score normalized population activity surrounding the lever press revealed distinct, phase-specific temporal profiles within the IL. To isolate core computational signals, we refined the analysis by excluding neurons that showed no significant modulation from baseline and labelled the remaining neurons as “tuned neurons”. We further divided these tuned neurons into significantly activated and significantly suppressed neuronal subsets.

During self-administration, average neuronal activity gradually decreased leading up to the lever press, forming a trough at the moment of the action (-0.76z, 50ms after LP), followed by a sharp, transient post-press returning to baseline (**Fig. 4A1**). While tuned neurons exhibited a similar trajectory, they were characterized by a shallower LP trough (-0.86z, at the LP) and higher post-press activity (0.22z, after 1.45s of lever press) (**Fig. 4A2**). These dynamics were driven by a robust contribution from activated neurons having an increased activity after the lever press (0.86z, 1.55s after LP) (**Fig. 4A3**) and a more sustained muted response from suppressed neurons (-0.82z, from LP to 3.85s after LP) (**Fig. 4A4**), potentially reflecting a neuronal signal that builds anticipation or motor preparation followed by a representation of the action’s execution or consequence.

**Figure 4:**
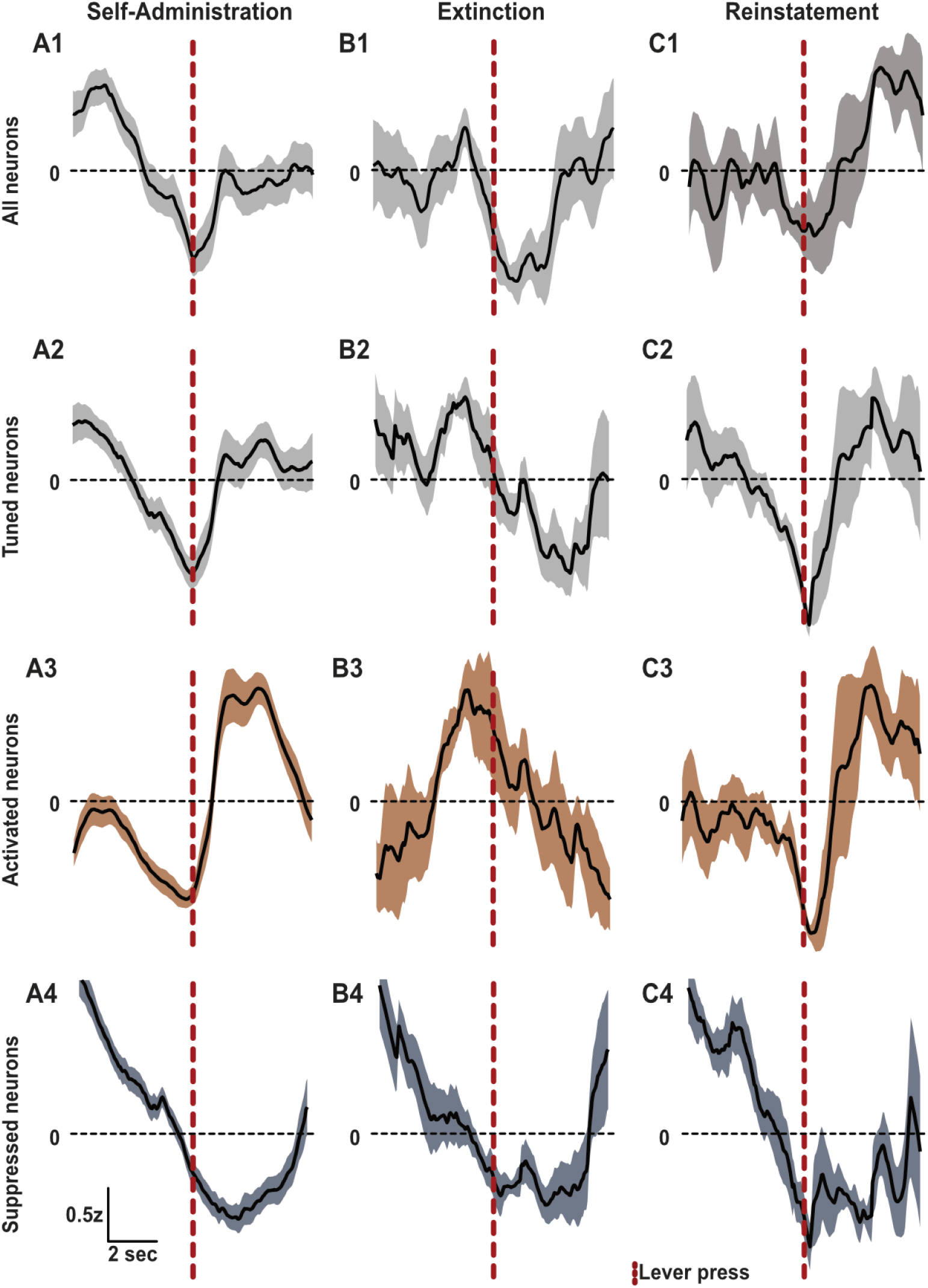
Characteristic tuning curves for different population of cells across different session types. **A–C** Population-averaged z-scored activity aligned to lever press (red dashed line) during self-administration (**A**), extinction (**B**), and reinstatement (**C**). Within each stage, rows show activity for all neurons (**1**), tuned neurons (**2**), activated neurons (**3**), and suppressed neurons (**4**). Vertical red dashed line represents lever press.

In contrast, the extinction task condition was characterized by a marked shift in the IL activity profile. Population traces revealed a prominent peak (0.5z, 1.15s before LP) approximately 1 second prior to the LP, followed by a sustained decrease (-0.7z) neurons for up to 2 seconds post-press (**Figure 4B1**). This profile was mirrored in the tuned neuron population (**Fig. 4B2**), where the pre-emptive activation was driven by a surge in firing from activated cells (1.2z, 1s before LP) (**Fig. 4B3**) alongside a relative sustained suppression from suppressed cells (-0.5z, from LP to 3.5s after LP) (**Fig. 4B4**). This pre-emptive firing likely signifies the recruitment of an inhibitory process to suppress the previously learned response, while the subsequent prolonged inhibition reinforces the updated non-reward contingency.

The reinstatement phase displayed an activity profile largely consistent with initial learning, characterized by a pre-press decrease (-0.65z at LP) (Figure 4C1-2). A key difference, however, was a substantial, delayed increase in activity during the expected reward window (0.88z, 3s after LP). Further analysis revealed that this unique activity distinct from the self-administration profile was driven primarily by an ensemble of neurons recruited during both extinction and reinstatement (**Supplementary Fig. 2**). While these neurons showed high pre-press activity during extinction, they transitioned to a bimodal pattern during reinstatement, firing both before the lever press and during the expected reward window. This suggests these neurons may encode a prediction correction signal, potentially serving as a ‘STOP’ command to suppress the conditioned response. Furthermore, the pre-press firing of this subpopulation explains the altered geometry of the reinstatement profile, specifically the narrower and right-shifted appearance of the activity trough. It is also due to this additive effect of neurons activating immediately after a lever press, and the additional delayed activated neurons, we find the percentage of activated neurons significantly higher in reinstatement compared to other task conditions (**Fig. 3D**).

In summary, the IL cortex exhibits distinct neuronal signatures that track the evolving contingencies of each task condition. Self-administration is characterized by broad pre-action suppression followed by post-press firing that encodes the fundamental task structure. Extinction shifts these dynamics toward pre-emptive firing before the action. Finally, reinstatement represents a functional combination of these two states. Together, these findings position the IL as a dynamic arbiter that integrates motivational cues with updated correction signals which motivated us to investigate the dynamics, and profiles of individual ensembles across the task conditions.

### Dynamic reconfiguration and stabilization of IL cortex ensembles during self-administration

How does the structured condition-specific activity emerge? To examine this, we first tracked the activity state of individual neurons across sessions and task conditions. **Fig. 5A** illustrates the spatial organisation of individual neurons from Rat 1. When comparing the spatial maps, it is evident that the activity of individual neurons is not preserved across behavioural conditions. Neurons frequently transitioned between being not significantly active, suppressed, or activated across learning days. Some neurons were active at self-administration and re-appear in reinstatement, yet there is no obvious pattern of re-occurrence of individual neurons among the three task conditions. This lack of a stable, topographical map suggests that the IL cortex does not rely on dedicated, stable ensembles of neurons to encode the learned action, rather it relies on representational drift models to encode the outcomes. This cellular dynamism is further highlighted by the state transitions between sessions, as shown in the Sankey diagram (**Fig. 5B)**.

**Figure 5:**
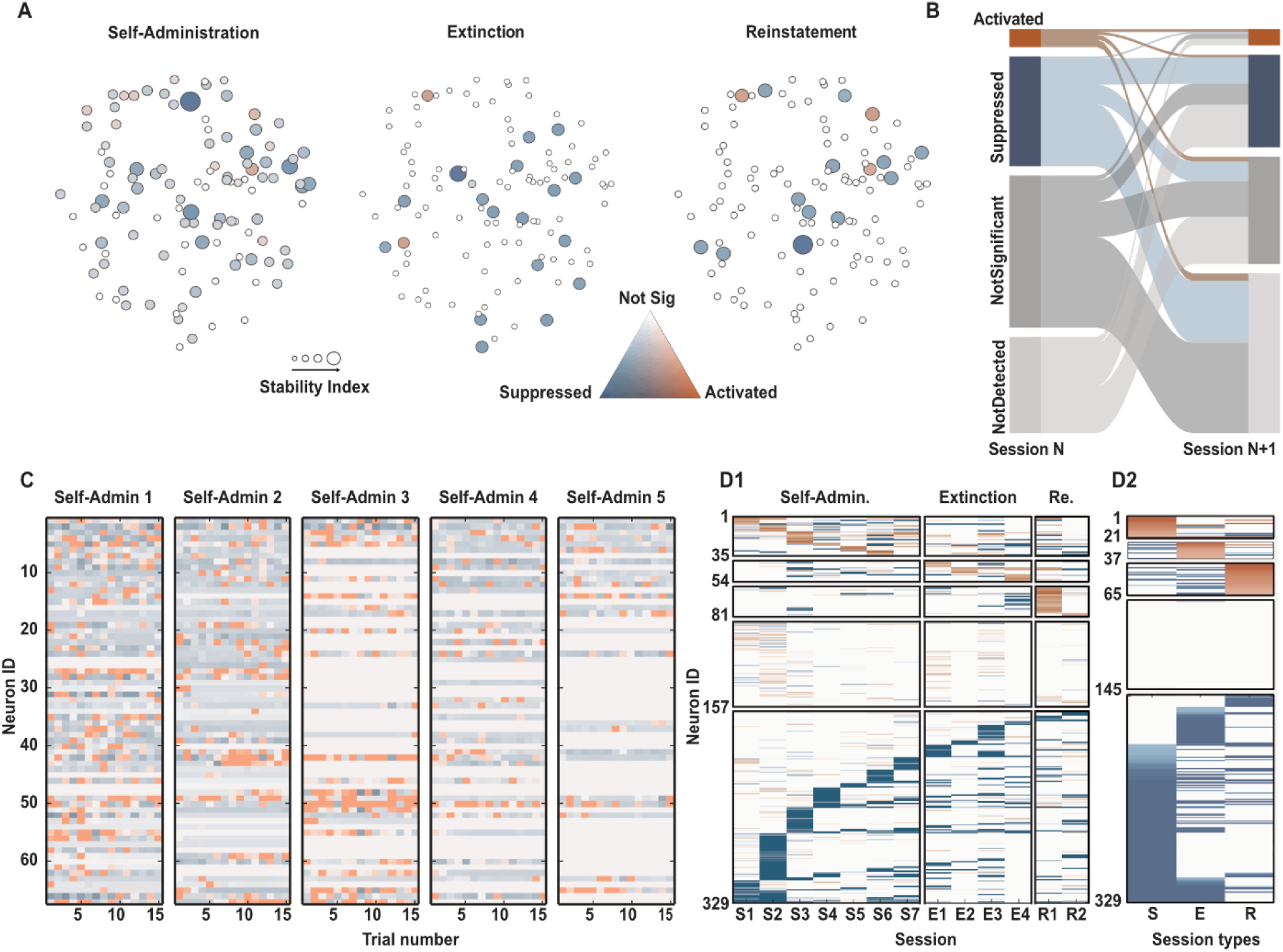
Dynamics of the neurons. **A.** Spatial footprints of recorded neurons from an example Rat 1, color-coded by their functional modulation: activated (orange), suppressed (blue), bi-directionally modulated (gray), or non-significant (white) relative to task events. The size of the cell represents its stability over the sessions of that type. **B.** Sander plot illustrating the transitions in functional cell identity between consecutive sessions (N to N+1). **C.** Trial-by-trial modulation heatmap across all neurons (y-axis) for first 5 self-administration sessions from Rat 1. Each cell is categorized per trial as excited, suppressed, or non-significant using the colour scheme in (**A**). **D.** All neurons from 3 rats (329 neurons) sorted hierarchically based on their activity across sessions (**D1**) and session types (**D2**), using the colour scheme in (A).

While individual neurons were unstable across days, the overall population activity showed a clear pattern of stabilization within and across sessions. Early in training, the tuning of the neuronal population appeared noisy and chaotic, but it settled into a more consistent and stable pattern in later sessions (**Figure 5C**).

To identify neuronal ensembles within this heterogeneous population, we analysed a subset of 329 neurons from the three rats that were tracked across all three task conditions. Neurons were sorted hierarchically based on the session in which they first exhibited significant modulation **(Fig. 5D)**. This organization revealed distinct functional clusters: the first three rows represent ensembles predominantly active during self-administration, extinction, and reinstatement, respectively. The remaining rows categorize neurons that were either non-responsive (Row 4) or significantly suppressed (Row 5). As shown in **Fig. 5D2**, grouping neurons by their longitudinal activity profiles clearly delineates ensembles specific to each task condition.

### Entropy reduction reveals progressive stabilization of IL population dynamics

Neuronal dynamics were further quantified using entropy, a measure of system disorder or unpredictability. We found that entropy consistently decreased both across trials within a single session (intra-session) and across the first four sessions of self-administration (inter-session) (**Fig. 6A, B**). This decrease signifies that the neuronal ensemble activity becomes more structured, predictable, and refined with practice. Corroborating this, we observed that raw population activity showed high variance during the first 50 lever presses of a session (**Figure 6C1),** but adopted a more stable, characteristic curve during the subsequent 50 presses (**Figure 6C2**). Furthermore, the z-scored activity of significantly activated neurons across the first four self-administration sessions revealed the dynamic evolution of the characteristic activity curve of the ensemble as sessions progress (**Fig. 6D)**.

**Figure 6:**
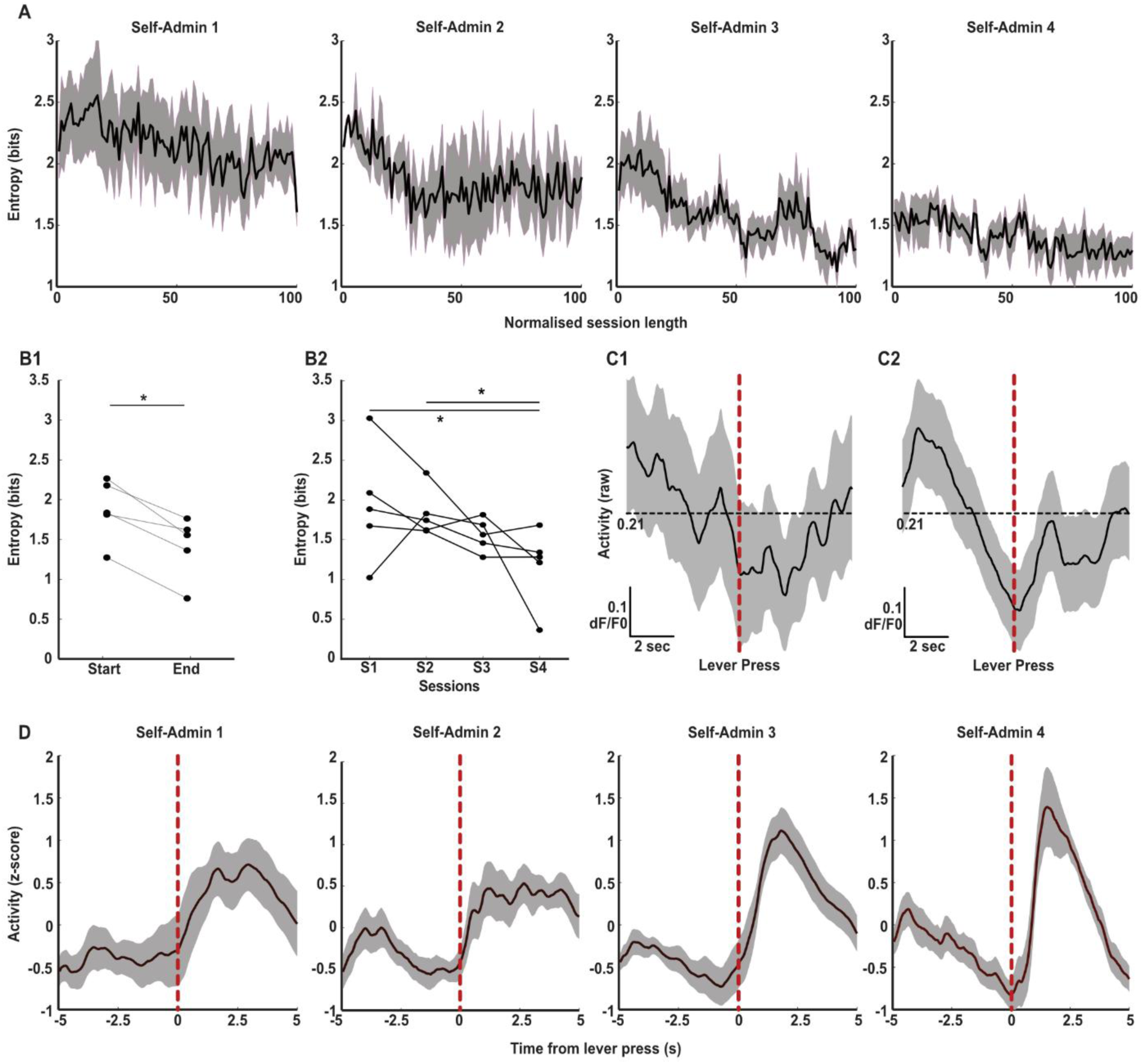
Stabilisation of the neural dynamics. **A.** Entropy as a function of normalized session duration (100 bins) for the initial five SA sessions (n = 5 rats). Solid lines indicate the mean; shaded areas represent ± SEM. **B-C.** Quantification of intra-session (B1) and inter-session (B2) entropy shifts across the self-administration stages. **C.** Comparative population dynamics during the first 50 lever presses (C1) versus the subsequent 50 lever presses (C2, trials 51–100). **D.** Z-scored activity of the significantly activated neurons across the first 4 self-administration sessions (n=5). *p<0.05, Wilcoxon’s signrank, LME.

To quantify this stabilization, we performed a Principal Component Analysis (PCA) on the population activity (**Supplementary Fig. 4**). Population activity during the initial trials of a session is highly scattered, indicating high variance, as the trials progress, the data points converge and cluster together demonstrating a reduction in variance. This suggests the neuronal ensemble begins each session in a high-variability, exploratory state and then settles into a stable and more efficient attractor state representing the well-rehearsed action sequence.

Together, these findings indicate that IL cortex implements a dynamic population coding strategy in which ensemble membership is highly flexible, while the population-level representation becomes progressively structured and predictable with learning. This stabilization suggests the emergence of an internal model of task contingencies. The dissociation between cellular instability and population-level stability is consistent with computational frameworks in which stable, low-dimensional representations arise from the dynamic recruitment of distributed neuronal subpopulations, enabling robust encoding despite heterogeneous single-cell activity ^37,38^.

### A predictive coding modelling of IL ensemble dynamics

To investigate a potential circuit mechanism capable of producing the observed neuronal dynamics in the IL cortex, we developed a computational simulation grounded in the principles of predictive coding and action-outcome learning ^30,31^. The simulation posits the existence of two functionally distinct neuronal populations, hereafter referred to as Ensemble 1 (E1) and Ensemble 2 (E2) which together learn and update internal representations of environmental contingencies (see Methods for a detailed description). This minimal architecture was designed to test whether interactions between predictive and error correcting populations are sufficient to reproduce the principal features of the experimentally observed IL activity patterns.

Ensemble 1 is conceptualized as the predictive ensemble consisting of 100 neurons with intrinsic temporal tuning distributed across the 10s trial window. Neuronal activity profiles were modelled using gamma functions parameterized to approximate the kinetics of GCaMP calcium transients. A Hebbian-like learning rule governs the plasticity of this ensemble: a successful prediction, where a neuron’s temporal tuning aligns with the occurrence of a salient stimulus (i.e., cue or reward), results in the potentiation of its response amplitude in subsequent trials proportional to its prediction accuracy. In addition, active E1 neurons exerted lateral inhibitory influences onto other E1 neurons proportional to their activity level, thereby promoting competitive population dynamics and sparse representations ^36,39^.

Ensemble 2 functions as the prediction correction ensemble. Consistent with theoretical frame proposing prediction-correction computations within medial prefrontal cortex ^30,31^,E2 activity increased selectively when expected outcomes failed to occur. Importantly, E2 activity reflected violations of predicted outcomes rather than stimulus salience itself. The magnitude of E2 activation scaled proportionally with the predictive strength generated by E1, thereby implementing a negative prediction-correction signal capable of suppressing outdated predictions through inhibitory feedback onto E1

Fig. 7A shows the activity curves of the E1 and E2 ensembles across training, extinction, and reinstatement trials. During self-administration neurons within E1 whose temporal tuning aligned with cue or reward delivery increased their activity across trials. Concurrently, lateral inhibition generated widespread suppression across the remaining neuronal population, reproducing the dominant inhibitory component observed experimentally during peri-event population analyses. Early training trials were characterized by weak predictive strength and correspondingly large positive prediction errors, whereas prediction errors progressively diminished as the model converged toward stable cue–reward representations

**Figure 7:**
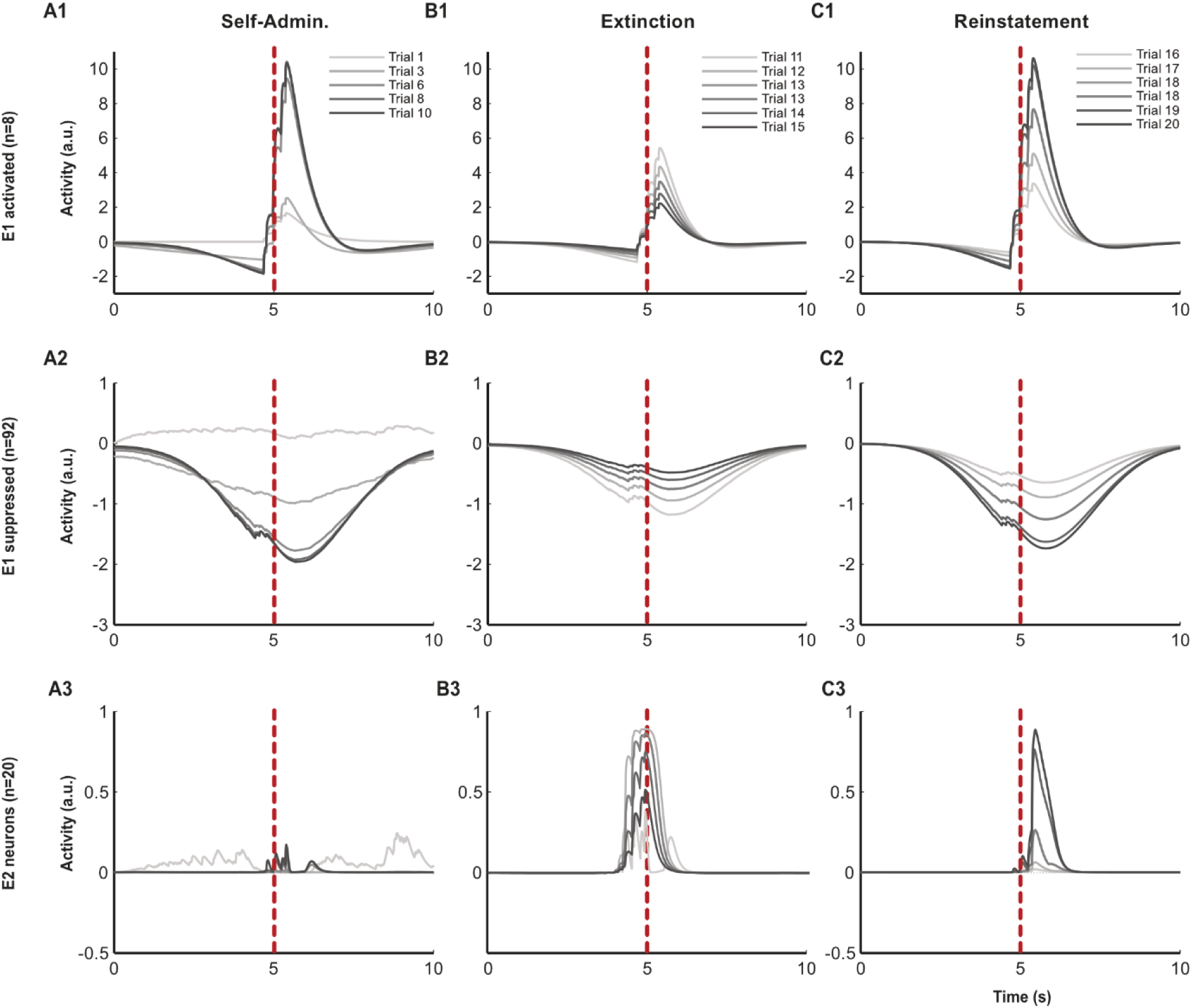
Dynamics of the simulation. Activity of simulated E1 and E2 neurons, over various trials. Trial 1-10: Self-Administration, Trial 11-15: Extinction, Trial 16-20: Reinstatement. **AX:** Self-Administration trials, **BX:** Extinction trials, **CX:** Reinstatement trials. **X1:** E1 activated neurons, **X2:** E1 suppressed neurons, **X3:** E2 neurons. Red dashed line denotes lever press.

During extinction, E1 initially continue to generate predictions for cue and reward delivery despite their omission. This mismatch produced strong negative prediction-error, hence stronger correction signals within E2 particularly around the expected reward period (**Supplementary Fig. 5**). Persistent omission of expected outcomes progressively weakened predictive activity within E1, resulting in reduced prediction strength across trials and the emergence of prominent pre-action activity patterns resembling those observed experimentally.

During reinstatement, reintroduction of the cue restored predictive activity within E1 associated with cue expectation. However, because reward delivery remained absent, E2 continued to generate negative prediction correction signals at the expected reward time point. This produced sustained post-action E2 activity closely resembling the delayed reinstatement-related activity observed in the empirical recordings. We hypothesize that persistent E2 activation contributes to the gradual suppression of reinstated responding by signalling continued violation of expected reward outcomes.

Importantly, the model also reproduced the progressive reduction in Shannon entropy observed experimentally during self-administration (Supplementary Fig. 4D). Entropy of simulated ensemble activity decreased across trials as population responses became increasingly structured and predictable. This reduction reflects the emergence of a more stable internal predictive model and parallels the experimentally observed transition from variable early-session dynamics toward low-variance population states.

Together, these findings demonstrate that a minimal predictive coding architecture based on interactions between temporally tuned predictive neurons and error correction neurons is sufficient to recapitulate the principal features of IL ensemble activity observed experimentally, including sparse excitation, widespread suppression, entropy reduction, and condition-specific peri-event dynamics. The model therefore provides a parsimonious mechanistic framework linking dynamic IL population activity to predictive representations of action–outcome contingencies and their updating during behavioural adaptation.

### Decoding behavioural states and task epochs from IL ensemble activity

To formally assess whether the structured population dynamics observed above encode task-relevant information, we applied supervised machine learning approaches to decode behavioural context and event timing from IL ensemble activity.

We first tested whether neural population activity discriminates between task conditions. 30 Fine-Gaussian Support Vector Machine (SVM) models were trained to classify 10 seconds (5 seconds before and 5 seconds after a LP, 201 frames) of z-scored traces from the three different task conditions, self-administration, extinction, and reinstatement. All the models were able to classify the data with high accuracy (Training: 96.22%, Test: 89.55%) and showed significant differences from models trained on shuffled data (Accuracy: 34.39%), demonstrating that the activity of IL cortex shows significant differences between different task conditions. (p=2.8e-06, **Fig 8A, B**)

**Figure 8.**
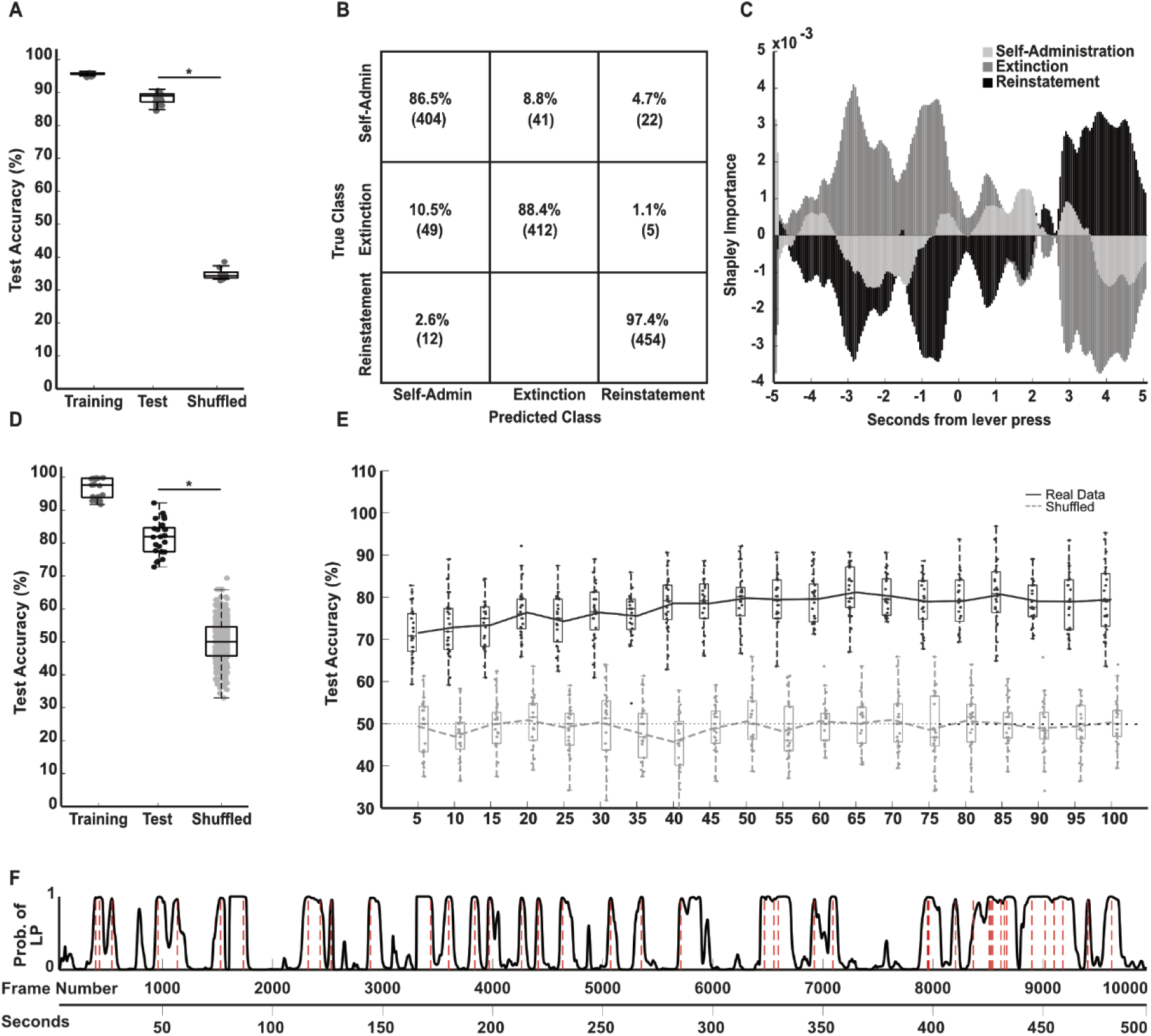
Machine learning-based classification and prediction of behavioural states. **A.** Training, test, and shuffled accuracies for the SVM models trained (n=30). **B.** Confusion matrix for the SVM classifier illustrating the accuracy of session-type predictions across the task conditions. The numbers in the bracket show the number of trials classified in the respective class. **C.** Temporal SHAP (Shapley Additive Explanations) importance curves for the three task conditions, illustrating feature contribution to the model as a function of time relative to the lever press. **C.** Comparative performance (training, test, and shuffled accuracy) for the Neural Network (NN) models trained with best-performing parameters (See methods, 5 seeds x 5 rats, 25 models). **E.** Test accuracies of NN models trained with the best-performing parameters on real (black) vs shuffled (grey) data. **F.** Probability of a trial occurrence quantified by the sample NN model (seed = 1) on a representative session. (Rat 1, S1). Red dashed lines indicate a rewarded lever press. *p<0.05, Wilcoxon’s signrank test.

To understand what features the model was using for this classification, and that there is no overtraining, we conducted a Shapley (SHAP) analysis (**Fig. 8C**). This revealed that the model’s predictions relied heavily on specific time windows within the trial and not just single instances. Crucially, frames just before the lever press were most important for identifying extinction trials, while frames approximately 3 seconds after the lever press were key for identifying reinstatement trials. This aligns perfectly well with our earlier findings of increased neuronal activity during these specific epochs, confirming that the SVM learned to leverage these biologically relevant time periods.

We next investigated whether IL activity encodes the temporal occurrence of task-relevant events. To test this, we trained feedforward neural network (FFNN) classifiers (MATLAB fitcnet) to distinguish “trial” activity, defined by a fixed frame window (F) around Lever Press (LP) events from “non-trial” control segments for each of the five rats. Control windows were selected using a distance-maximization algorithm to ensure they were temporally isolated from LPs while maintaining identical dimensions. For each event, calcium traces from N neurons were flattened into a N x F feature vector and standardized for classification.

To ensure the reliability and generalizability of the decoder, we performed an exhaustive parameter search, training 4,000 model iterations across various architectures, regularization parameters, and preprocessing steps (Methods). The classification performance of all the models was robust across the entire parametric space. (Training Accuracy: 99.8%, Test Accuracy: 75.58%, **Supplementary Fig. 5**).

Using an ANOVA to compare model performances, we identified that a 25-layer architecture (p = 1.2e-07), the exclusion of PCA (p = 1.4e-17), and a 90/10% train-test split (p = 1.1e-04) yielded significantly higher test accuracies. While the inclusion of a regularization parameter (λ = 0.05) improved performance specifically for larger temporal windows (p = 5.7e-04), data augmentation was found to increase training accuracy without improving test performance (p = 0.92; **Supplementary Fig. 5)**. To confirm that the models captured genuine biological insights rather than artifacts of data processing, we conducted permutation testing. Using the best-performing parameters, we trained 500 control models (100 permutations per rat) on shuffled labels. These control models performed near chance levels, while the original models achieved significantly higher accuracy (81.9% vs 50%, p = 2.98e-17, **Fig 8D, Supplementary Fig 5**).

To identify the optimal temporal scale required to classify trial information from IL activity we systematically varied the input duration from 5 frames (0.25s) around an LP to 100 frames (5s), we were able to train models which performed significantly above the shuffled controls for all the windows (p=0, **Fig 8E**). We found that test accuracy saturated at a window of 50 frames (±2.5s). Finally, applying these trained models in a sliding-window approach across continuous recordings revealed that prediction probabilities peaked near trial events, reliably detecting the occurrences of the trials (**Fig. 8F**).

Together, these results demonstrate that IL population activity contains decodable signatures of both behavioural context and event timing. This supports the view that the progressively structured and low-entropy population dynamics observed during learning correspond to functionally meaningful representations that can be read out by downstream systems.

These constraints argue for a circuit architecture in which temporally structured predictive representations are continuously shaped by error correction driven inhibitory feedback, supporting our hypothesis formalized in the dual-ensemble model described above. This shows that the models learned the specific ensemble activity patterns associated with the core task event. Together, these results highlight that IL population activity is not just randomly dynamic, but contains a rich, decodable structure that reliably represents both the broader behavioural context and the precise timing of specific actions within a task.

## Discussion

The infralimbic cortex (IL) has historically been interpreted either as a behavioural “STOP” system mediating extinction or as a facilitatory region supporting reward seeking and habit formation. Our findings suggest that these apparently opposing functions can be reconciled. Longitudinal calcium imaging across acquisition, extinction, and reinstatement showed that IL activity is organized into sparse, dynamically evolving ensembles whose temporal structure is tightly linked to behavioural contingencies. Rather than acting as a simple inhibitory switch, the IL appears to implement predictive representations of action outcomes together with prediction correction-related signals that update these representations when contingencies change.

Previous lesion, optogenetic, and immediate early gene studies established that distinct IL ensembles contribute to extinction, reward seeking, and habit-like behaviour ^3,18,40–42^. However, these approaches generally lacked the temporal resolution necessary to resolve how ensemble activity evolves during behaviour and across learning stages. Our longitudinal recordings revealed that only a small fraction of neurons exhibited significant event-related activation, whereas a substantially larger fraction showed suppression during task execution. This sparse coding regime embedded within widespread suppression resembles observations in electrophysiological studies of IL and orbitofrontal cortex during learned behaviour ^25,41^ and is consistent with theoretical frameworks proposing that predictive representations are sharpened through suppression of non-informative population activity ^35,36^.

Importantly, the temporal organization of these ensembles differed systematically across task conditions. During self-administration, population activity exhibited a pronounced decrease preceding the lever press followed by a transient post-action increase, consistent with a predictive representation of expected outcomes. In contrast, extinction was characterized by strong pre-action activation followed by sustained suppression. Within our framework, this pre-action activity may reflect recruitment of a prediction correction ensemble that becomes engaged when expected outcomes fail to occur. This interpretation provides a mechanistic explanation for why IL disruption impairs extinction learning in many behavioural paradigms ^6,33^. Rather than directly suppressing behaviour per se, IL activity may encode violations of predicted outcomes that drive behavioural updating. Reinstatement further clarified this dual functionality. Reintroduction of the cue restored reward expectancy and increased behavioural engagement despite continued reward omission. At the neural level, reinstatement partially recovered the activity structure observed during acquisition but additionally recruited a delayed post-action component associated with increased proportions of activated neurons. This finding may reflect the increased computational demands imposed by reinstatement, where previously learned contingencies become ambiguous and require simultaneous maintenance of competing predictions. Such ambiguity-related recruitment is consistent with broader theories assigning medial prefrontal cortex a role in conflict monitoring and adaptive control ^1,30^. In the context of addiction, this increased computational demand may be particularly relevant, as impaired predictive updating or excessive weighting of cue-related predictions could contribute to relapse vulnerability.^43^

A central finding of the present study is the dissociation between instability at the level of single neurons and stability at the population level. Individual neurons frequently changed their tuning properties across sessions, yet ensemble-level dynamics became progressively more structured and predictable with learning. This stabilization was accompanied by a robust reduction in Shannon entropy both within and across sessions, indicating convergence toward lower-dimensional population representations. Such dynamics are compatible with models in which cortical circuits initially explore a high-dimensional representational space before converging onto stable attractor-like states that support efficient behavioural policies ^37,38^. Notably, these observations also provide a potential bridge between predictive coding and classical theories of habit formation, suggesting that habitual behaviour may emerge from progressively compressed and stabilized predictive representations. From a translational perspective, the presence of functionally distinct predictive and error-related ensembles suggests that successful neuromodulation of cognitive flexibility may depend less on the targeted brain region than on the specific neuronal populations recruited within it.

To formalize these observations, we implemented a minimal predictive coding model comprising temporally tuned predictive units and prediction-correction units. The model reproduced several core empirical findings, including sparse tuning, widespread suppression, entropy reduction, and condition-specific temporal dynamics. Importantly, machine learning analyses demonstrated that IL population activity contains highly decodable information about both behavioural context and event timing, with classifiers relying specifically on the temporally structured epochs identified experimentally. While these findings are consistent with predictive coding frameworks, alternative interpretations based on salience encoding, action monitoring, or state-dependent inhibitory control cannot be excluded. The present model should therefore be viewed as a parsimonious computational account rather than a definitive circuit reconstruction.

Several limitations should be considered. The number of animals completing the full longitudinal paradigm was limited, reflecting the substantial technical challenges associated with chronic deep brain miniscope imaging in freely moving rats. Maintaining stable optical access across months of operant training, extinction, and reinstatement represents a major experimental constraint that currently limits throughput relative to mouse imaging approaches. Nevertheless, ensemble dynamics were highly consistent across animals and conditions. In addition, our conclusions remain correlational, and future experiments combining longitudinal imaging with ensemble-specific manipulations will be required to establish causal contributions of predictive and correction-related populations.

Together, these findings support a reinterpretation of IL cortex function as a dynamic predictive system that continuously updates behavioural representations in response to changing contingencies. By linking sparse ensemble coding, entropy reduction, and prediction dynamics within a unified longitudinal framework, the present study provides a mechanistic account capable of reconciling previously conflicting views of infralimbic function.

## Conclusion

In conclusion, our findings suggest a fundamental shift in how we conceptualize the infralimbic cortex, moving away from the traditional view of a simple inhibitory switch toward a more nuanced role as a predictive engine. By identifying the specific dynamics of dual ensembles through miniscope imaging, we have provided a mechanistic explanation for the previously irreconcilable “go” and “stop” functions. The observed reduction in entropy and the emergence of a refined 10% task-responsive population during acquisition align with the region’s established role in habit formation, while the activation of pre-press correction signals provides a clear neuronal basis for the process of extinction. This framework not only synthesizes decades of conflicting research but also offers a predictive model that accurately identifies behavioural states based on real-time neuronal dynamics. Ultimately, the IL cortex serves as a critical node for updating behavioural strategies, ensuring that an organism’s internal predictions remain accurately aligned with environmental realities.

## Methods

### Animals

Wistar rats were ordered from Charles River and kept at an inverted 12-hour dark/light cycle with access to food and water ad libitum. At the start of the experiments animals were 8 weeks old. All procedures were approved by the local animal welfare body (Regierungspräsidium Karlsruhe, G-221/15, In-vivo Mikroendoskopie im Gehirn von Ratten). Data from five rats was analysed, they have been labelled as Rat 1, Rat 2, Rat 3, Rat 4, and Rat 5 across the paper.

### Miniature UCLA microscopes

UCLA Miniscopes were manufactured based on the descriptions of the miniscope.org webpage. The CMOS imaging printed circuit boards (PCB) and the data acquisition (DAQ) PCBs were obtained from SierraSircuits based on the designs the developers released on the GitHub repository (PCB and DAQ version 3.2 (Aharoni, 2016). Later additional CMOS and DAQ PCBs were bought from labmaker (Berlin, Germany). The housing (focus slider, main body and filter cover, all black Delrin) as well as the baseplates (aluminium) and the baseplate-covers (black Delrin) were manufactured by the local machine-workshop and were also based on the UCLA Miniscope design. Achromatic and half-ball lenses were bought from Edmund optics (5×15 mm achromatic lens, ID:45207; 5×12.5 mm achromatic lens, ID:49923; 5×7.5 mm achromatic lens, ID:45407; half ball lens 3 mm, 47269). GRIN lenses were bought from Inscopix. The circuit boards for connecting the emission diode and the coaxial cable connector were obtained from SilverCircuits (Houston, USA) and based on the designs the developers released on the GitHub repository (Aharoni, 2016g). The emission LEDs and miscellaneous electronics and items were bought from Digikey (Thief River Falls, USA; LED Luxeon Rebel Blue SMD, ID: 1416 1028 1 ND; SMA Connector 50 Ohm, ID: consma013.062; Light Pipe clear 3mm, ID: VLP 550 F; IC Eeprom 1Mbit 400Khz, ID: 24AA1025 I/P; shunt jumper .1”, ID: 969102 0000 DA; IC DIP socket 8pos, ID: DILB8P 223TLF; SMA jack R/A 50 Ohm, ID: consma002). Optic filters were bought from Chroma (Bellows Falls, USA; excitation filter, ET470/40x 3.5×4×1mm, ID: IN054535; emission filter, ET525/50x 4×4×1mm, ID: IN054538; dichroic mirror, T495lpxr 4×6×1mm, ID: IN054536). Magnets for the baseplates and main bodies were purchased from KJ Magnetics (Pipersville, USA; 1/16” dia. x 1/32” thick, ID: D101-N52). The housing for the DAQ boards was 3D printed on an Ultimaker 2 using 3 mm thick, white PLA at 100µm z-resolution (Aharoni, 2016). Two different laptops (Dell Inspiron 15, MacBook Pro 15, 2017) were used for the recordings. Both had windows 7 installed as the operating system and the recording software supplied by the UCLA Miniscope team was used for calcium imaging.

### Surgical Procedures

#### Viral injection and GRIN lens implantation

Animals were anesthetized with Isoflurane (1.5 - 3%, Univentor 1200 - anaesthesia Unit, Univentor, Zejtun, Malta) and head-fixed in a Kopf Stereotaxic frame (Model 900, David Kopf Instruments, Tujunga, USA). A small dose of dexamethasone was injected to reduce the intracranial pressure (reference). The head was shaved to expose skin and then disinfected with ethanol swabs. The skin was cut along the anterior-posterior axis and the connective tissue was removed using cotton swabs to expose the bone. Three small craniectomies were drilled using a dental drill and Fine Science Tools (FST, Foster City, USA) bone screws were fixed to the skull. A craniectomy (∼2mmØ) was made at 3mm AP, and 0.5mm ML to bregma. A pulled glass injection pipette was used to inject AAV1.Syn.GCaMP6f.WPRE.SV40 (UPenn Vector Core, Philadelphia, USA, Lot#: CS1107, titer: 2.13*1013 GC/ml, 1:20 dilution in PBS, 500nl) at 5mm depth. 500 nl of the virus was then slowly injected at the speed of 100 nl/min. Before the insertion of the GRIN lens, a glass fiber (Ø 127µm) was slowly lowered upto the depth of 4.4 mm from the surface of the skull to create a path for the GRIN lens. The GRIN lens (1 mm x 9 mm, Inscopix, Palo Alto) was lowered to the depth of 4.7 mm into the brain. A modified Luigs&Neumann linear actuator was used to lower the injection needle, glass fiber, and the GRIN lens into the brain. The linear actuator repeatedly lowered the object 300 µm below and then raised the object 200 µm above to alleviate pressure. The alternation rate was changed to lowering of 30 µm followed by raises of 20 µm, as the objected reached 1 mm from the target site. After the insertion of the GRIN lens, it was fixed using a layer of OptiBond™ FL Kit (Kerr, Bioggio, Switzerland) followed by dental cement liquid (Paladur, Henry Schein, Gallin, Germany) and black dental powder (Contemporary Ortho-Jet™, black, Lang Dental, Wheeling USA).

#### Baseplate Implantation

Baseplates for the UCLA miniscope system were equipped with magnets and a set screw to allow a tight connection to the miniaturized epifluorescence microscopes. Before the microscope was attached to the stereotax, the focus was centered to its mid-range to allow adjustments in both directions after the fixation of the baseplate. Animals were anesthetized with isofluorane (1.5 – 3%, Univentor 1200, anaesthesia Unit, Univentor, Zejtun, Malta) and head-fixed in a stereotaxic alignment system (Kopf Stereotax, model 900, Tujunga, USA). The protective cement cover of the implanted GRIN lens was carefully removed using a dental drill and the lens paper in the cement cavity was carefully removed with forceps. Care has to be taken to avoid scratching the surface of the GRIN lens. After the surrounding cement was drilled down to the level of the lens and all the lens paper was removed, the lens was cleaned with lens paper and ethanol. Care was taken to 23 remove all remaining vacuum grease. In case of contaminations of the lens with dental cement, acetone was used to dissolve the cement and clean the surface of the lens. Care has to be taken to not put acetone in contact with the wound or the skin of the animal. Once the lens was clean and could be reached with the objective GRIN lens, the miniaturized microscope, attached to the stereotaxic frame by a custom built holder, with the attached baseplate was lowered down to the implanted GRIN lens. The miniaturized microscope was attached to a PC running Windows 7 (Dell, Inspiron 15) through the DAQ board and the software supplied by the developers (Aharoni, 2016f) was running to provide visual feedback on the field of view (FOV) of the microscope. Once the GRIN lens was within the focal distance of the microscope the imaging parameters were initially adjusted to optimize the range of the histogram and facilitate the process of focusing. The image repetition rate was generally set to 20 Hz (corresponding to exposure times of approximately 50 ms per frame), except for cases of weak calcium signals. The gain was always set to the maximum value and the LED power was adjusted until the background was visible, but no over-saturation occurred. Because of the anaesthesia, it was rarely possible to see calcium transients of individual cells. To find an appropriate field of view for the recordings, the bottom of the GRIN lens was brought into focus (close to the top of the GRIN lens). From there the microscope was raised until no more scarring on the bottom of the lens was visible, but there were still blood vessels in focus. If a satisfying focal plane was identified, dental cement liquid (Paladur, Kulzer Mitsui Chemicals Group, Hanau, Germany) and black dental powder (Contemporary Ortho-Jet™, black, Lang Dental, Wheeling USA) were mixed and the four corners of the baseplate were attached to the implant with the fresh cement. Care was taken to avoid using cement that was too liquid to avoid the cement ‘running’ below the baseplate and covering the lenses. After the four corners of the baseplate were attached to the implant, the remaining sides were covered in cement. To avoid detachment of the aluminium baseplates from the cement, super glue (Pattex, flüssig, Düsseldorf, Germany) was carefully applied to the edges between the aluminium baseplates and the dental cement. Care was taken to avoid getting super glue between the baseplate and the microscope to avoid permanent attachment of the two. After the baseplate was cemented in place, the microscope set screw was carefully opened and the microscope could be carefully removed. The baseplate was then covered with a baseplate cover (Aharoni, 2016g) (UCLA, Zentralbereich Neuenheimer Feld, machine shop) to protect the lens from dirt.

### Behavioural procedures

#### Operant self-administration hardware

All experiments were performed in operant chambers (MED Associates chamber, ID: MED-008- CT-B4) enclosed in ventilated sound-attenuating cubicles. The chambers were equipped with a response lever on each side of the chamber. Responses at the appropriate lever activated a dipper cup in which the reward was presented inside of a head entry port. A light stimulus was placed above the active response lever of the self-administration chamber. An IBM-compatible computer controlled the delivery of fluids, presentation of stimuli, and data recording. TTL pulses from the operant chamber for the levers and head entry ports on both sides, as well as the cue light for the rewarded side were acquired/digitized using an Arduino Mega running the Firmata (Firmata, 2015) firmware, along with the frame trigger of the DAQ board of the miniaturized microscope. The data that was received by the Arduino was then saved using Bonsai (Lopes et al., 2015) as .csv file.

The conditioning chamber was 3D modelled in Blender 2.9.

### Operant conditioning

#### Self-Administration

Saccharin self-administration training and testing sessions were performed 3 h after beginning of the dark phase, 5–6 days per week. Animals were trained to self-administer 0.2% (w/v) saccharin in daily 30 min sessions on a fixed-ratio 1 schedule using a saccharin fading procedure modified from Samson, Pfeffer, & Tolliver, 1988.^44^ During the first 3 days of training, the animals were kept water deprived for 20 hours per day. Responses at the active lever were reinforced by the delivery of 0.2% (w/v) saccharin solution. For the remaining experiments the animals underwent the same procedure without water deprivation. Responses at the inactive lever were recorded but had no consequences. A (visual) cue was used for conditioning. The discrete visual stimulus was presented after correct responses resulting in saccharin delivery (active lever). As visual stimulus, a 5 s blinking light was used, which was activated after a response at the active lever and was therefore directly connected to saccharin availability. The 5 s period served as a “time out”, during which responses were recorded, but did not lead to an additional reward delivery (conditioned stimulus, 1st 1.5s, 2nd 1s, 3rd 1s; 1s off in between pulses). For the stimulus-conditioning training, the animals had to complete 10 sessions of 30 minutes, with the conditioned stimulus (CS), without simultaneous calcium imaging, and after baseplate fixation enough sessions to reach previous lever response levels with simultaneous calcium imaging. A Med Associates Super Port Input module (ID: DIG-712) was used to relay TTL pulses indicating behaviourally relevant events in the chamber (lever press left/right, head entry left/right, cue 25 light left/right). Upon pressing the rewarded lever (always the right side), the dipper cup was placed in the liquid and the reward was presented in the head entry port. The pre-training was done by Simone Pfarr, Janet Barroso-Flores, Laura Schaaf, and Rebecca Hoffmann.

#### Extinction

After successful completion of the stimulus conditioning phase all animals underwent 4 daily, 30 min extinction sessions, which was sufficient to reach an extinction criterion of 25% of the baseline activity at the active lever per session. During extinction sessions, both levers were extended. Responses at the previously active lever activated the dipper cup, which did not result in reward delivery or presentation of the discrete CS (blinking light). The TTL-pulse was still transmitted to aid the post-hoc analysis.

#### Cue-induced reinstatement

For the cue-induced reinstatement sessions, both levers were extended during the sessions and the animals were presented with the same conditioned stimuli (CS) as during the conditioning phase. after pressing the rewarded lever. The dipper cup would move; however, no reward was presented in the head-entry port.

### Histology

The animals were deeply anaesthetized with isoflurane and transcardially perfused with 100 ml 1xPBS (137 mM NaCl, 2.7 mM KCl, 8 mM Na2HPO4, 1.46 mM KH2PO4, pH 7.4), containing 10000 units of Heparin sodium/l. Next, the animals were perfused with 50 ml of 4% paraformaldehyde (PFA) in 1xPBS solution (pH 7.4). The complete heads of the animals were post fixed in fixative solution for at least 7 days at 4 °C. After removal from the skull, brains were stored in 4 °C PBS. Perfusions were performed by Simone Pfarr and Janet Barroso-Flores.

### Thin sectioning and mounting

After at least one week of incubation in fixation solution at 4 °C the brains were dissected from the skulls. The GRIN lenses were removed along with the top of the skull before dissection. Care was taken to avoid damaging the tissue with the implanted GRIN lenses. After the dissection the 26 brains were rinsed 2x in fresh PBS. Then the brains were cut in half along the anterior-posterior axis to separate the hemispheres. The side which contained the GRIN lens was then cut a second time approximately 3 mm posterior to the entry site of the GRIN lens and the anterior half was mounted in a vibratome Slicer (Leica VT1000S) and cut into 100 µm thick sections. The sections were collected in 24 well plates containing PBS with DAPI (#D9542-10MG, final concentration 0.05-0.1 mg/ml, Sigma-Aldrich, St. Louis, USA). Those slices close to the area which contained the GRIN lens were mounted on 24 mm x 70 mm glass specimen slides (Marienfeld, LaudaKönigshofen, Germany), covered with SlowFade (#S36936, Thermo Fisher Scientific, Waltham, Massachusets) and a coverslip (24 mm x 60 mm, Carl Roth, Karlsruhe, Germany) and then sealed with nail polish.

### Image acquisition of fixed tissue sections

Overview images were acquired using a Leica DM6000 upright, fluorescence microscope, equipped with a 1.25X/0.04 NA (Leica Microsystems, #11506215) and a 10X/0.4 NA (Leica Microsystems, #11506284) objective, and using the LasX acquisition software. The emission light-path contained an additional magnification step of 1.2x to account for the size of the camera. High-resolution close-up images were acquired using a Leica SP8 confocal microscope, equipped with a 10X/0.4NA (Leica Microsystems, #11506293) objective, using the same software. The image resolution was kept at 1024×1024 pixels, while the scan speed was set to 200 Hz with solely unidirectional scanning. All fluorophores were imaged sequentially. Image processing was performed using Fiji (Rueden et al., 2017).

### Analysis

#### Analysis of operant conditioning data

The operant conditioning setup provided six data streams which were collected from Bonsai, namely cue-light, right and left levers, right and left head entry ports, and miniscope frame trigger. The TTL data from these streams was arranged in structures in MATLAB, and then analysed and plotted using MATLAB.

#### Calcium imaging data extraction

The raw videos were analysed Minian ^45^. The analysis was carried out on the Baden-Wüttemburg High Performance Cluster (bw-HPC) and on a local PC. The Xarray files from Minian were exportd into a combined ncdf (.nc) database after which they were imported into MATLAB, and arranged into structures to carry out further analysis. Minian analysis jupyter notebooks are provided in the supplementary data.

#### Alignment of calcium imaging and behavioural data

The sixth data stream with the miniscope frame trigger was used to align the behaviour data and the miniscope frames. In the case of frames dropped, the timestamps were looked at manually and were matched accordingly. All the timestamps from the operant condition data streams were converted to their respective frame numbers and were used for further analysis.

#### Matching cells between sessions

To match the cells across the sessions two independent pipelines namely, CellReg ^46^ and CrossReg ^45^ were used. Upon inspecting the performance of the two pipelines, CrossReg was chosen was all the further analysis. The data were then rearranged for the structures to always have the same cell on the same index throughout sessions. The neurons were then manually scanned through and discarded if they were too small or had irregular non-neuronal activity graphs. The number of cells detected, matched and discarded are shown in **Supplementary Figure 6.**

#### Removal of fluorescent decay

The extracted data were cropped to maximum of 30 mins to avoid any major fluorescence decay, and was further processed using an Asymmetric Least Squares (ALS) smoothing, to estimate and subtract the decay ^47^, using the formula:

The objective function used to find the baseline:

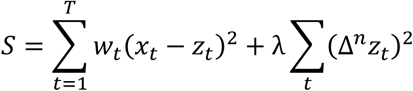

Where, *x_t_* is the original fluorescence trace at time *t*, *z_t_* is the estimated baseline, *w_t_* is the asymmetric weight at time t, *λ* is the smoothness penalty parameter (params.lambda, default: 10e-7), Δ*^n^* is the difference operator of the order *n,* (params.order, default: 2).

To minimize the above function the script solves the following linear equations at each iteration:

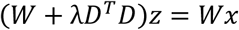

Where, ***x*** is the trace column vector, ***z*** is the baseline column vector, *W*is a diagonal matrix containing the weights *w_t_*, *D* is the matrix representation of the difference operator.

After each basline estimation the weights are updated according the the following:

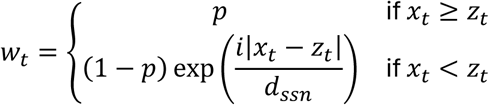

Where *p* us the asymmetry parameter (params.p, default: 0.05), *d_ssn_* is the sum of the absolute values of all negative residuals: *d_ssn_* = ∑*_xk_*_<*zk*_|*x_k_* − *z_k_*|

Finally, the baseline (decay thereof) is subtracted from the original trace to get the correct value.

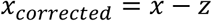

Example uncorrected and corrected traces, for Rat 1 first 4 sessions are shown in **Supplementary Figure 7**.

#### Extraction of trial events

100 frames before and after each rewarded right lever press during self-adminstration, and all right lever press in cases of extinction and reinstatement were extracted, assuming the frames-per-second (FPS) to be 20. We acknowledge, that it is possible that due to frame drop or lower recorded frame rate, these 100 frames might not exactly represent 5 second window, the error for each session was calculated was kept to a minimum of 50ms, as shown in Table X (AlignmentTable).

#### Extraction of trial-tuned cells

Z-scores during the trial were calculated for all the cells individually, by normalising them by the mean and standard deviation as follows:

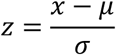

Where, z: z-scored value, x: raw trace value, µ: mean raw trace value, *σ*: standard deviation of the raw trace value.

These z-scores were then compared with the z-scores from the non-trial section bootstrapped 10,000 times. If a cell had a significantly different z-score during trial than a non-trial section, it was called trial-tuned cell. If z-score of a trial-tuned cell was higher than baseline, it was termed as a significantly activated neuron, if it was lower than baseline, it was termed as a significantly suppressed neuron.

### Entropy Analysis

Shannon’s entropy (S) for all the dynamics of the cells was calculated over a moving window of 100 frames. The activity was divided into 100 bins, and the entropy was defined as:

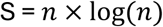

where n represents the number of bins of activity. For the simulation, no moving window was used, but the activity of E1 and E2 neurons was directly divided into 100 bins and the entropy was similarly calculated for each trial.

### Dual Ensemble Simulation model

A computational simulation was developed to model neural ensemble dynamics utilizing the Predicted Response Outcome (PRO) framework ^30^. The simulation evaluated the temporal activity profiles of two distinct neural populations: Ensemble 1 (E1), responsible for encoding temporal predictions of task events, and Ensemble 2 (E2), responsible for encoding surprise or prediction errors, and providing correction signals. The simulated paradigm was divided into 3 sequential phases, Self-Administration (where both cue and reward were present), Extinction (where no stimulus was present) and Reinstatement (where only cue was present).

The results shown in the paper are with using the default parameters, which are shown in brackets throughout the methods section.

The number of trials in each phase could be assigned by the parameter trialPhases ([10 5 5]), which takes in the number of trials to be simulated for each phase, self-administration, extinction, and reinstatement respectively. The total duration of each trial was defined by trialLength ([10]). The simulated time was vectorised into timepoints denoted by timeVec ([100] per trialLength, 1000 timepoints total), to get higher temporal resolution. Within each trial, the precise timing of the expected cue, cueTime ([5.3]), and reward, rewardTime ([5.7]), was subjected to a Gaussian temporal jitter defined by stimJitter ([0.01]) to simulate natural variability in stimulus delivery and perception.

The activity of each neuron was modelled as a Gamma curve (high rise, riseGamma ([0.3]), slow decay, decayGamma ([0.6])) and a preferential temporal tuning field center (µ) distributed evenly across the trial length and a baseline amplitude baseline_E1/baseline_E2 ([2]). The formula used to calculate the activity is as follows:

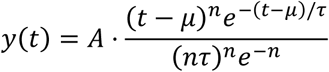

Where, t: current timepoint, µ: tuning center, A: peak amplitude, *n*: shape parameter (riseGamma), *τ*: time-scaling parameter (decayGamma).

The model simulated an ensemble of E1 neurons, numCells_E1 ([100]), divided into “cue-predicting” and “reward-predicting” cells based on a defined proportion, cueRewardRatio ([0.5]). During self-administration, the activity amplitude of E1 neurons scaled iteratively as a function of a set learningRate ([0.8]), up to a defined maximum threshold, maxActivity ([10]), depending on the overlap between the stimulus (cue or reward) and the activity of the E1 cell during the trial. To simulate local network competition, lateral inhibition, applyInhibition ([true]), was applied. A Gaussian-smoothed trace of the total E1 population activity smoothed by inhSigma ([1.5]) was calculated and subtracted from the uninhibited trace of each individual E1 neuron, scaled by a suppressionIntensity ([0.05]) parameter. The precise biological substrate for this inhibition, whether mediated by local neurons or top-down feedback from other cortical or subcortical regions, is not critical to the model’s core function.

Since the core of the simulation relies on mismatch between predictions (P_cue and P_reward) and the actual outcomes (E_cue and E_reward). The actual occurrences of the cue E_cue and reward E_reward were modelled as binary pulses smoothed by a Gaussian kernel. The population-level expectation for the cue (P_cue) and reward (P_reward) was computed as the mean positive activity of the respective E1 subpopulations.

Instantaneous prediction corrections, or “surprises” ω, were calculated at each time step. Negative surprise 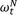, an expected event failing to occur and positive surprise 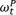, an unexpected event occurring were defined mathematically as:

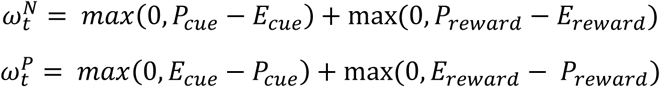

E2 neurons, numCells_E2 ([20]), were modelled to reflect the negative surprise trace 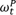 generated by the simulation, as a correction in the following trial. The instantaneous negative surprise was passed through a non-linear sigmoidal transfer function to map the mismatch value to corresponding E2 neural activation.

### Support Vector Machine Model

A fine gaussian support vector machine model was trained from MATLAB’s deep network toolbox’s classification learner. Data 100 frames before and 100 frames after a lever press (rewarded LP in case of Self-Administration) was extracted as a trial (total 201 frames, including LP). 30 models were trained with different seeds and were tested against 30 models trained with shuffled labels. To remove any class biases, the data for extinction and reinstatement was augmented by shifting the extracted data by +/- 1 frame. Further, Shapley Importance curves were plotted to demonstrate absence of overtraining or specific learning.

#### Neural Networks

For each of the 5 animals, neuronal data from Self-Administration sessions were aggregated by extracting a fixed frame window (F) around each Lever Press (LP) event. Non-LP control windows were selected using a distance-maximization algorithm to ensure they were temporally distant from any press and of the same size. For each event, calcium traces from N neurons were flattened into a N x F feature vector and standardized. We trained 4000 different Feedforward Neural Network (MATLAB fitcnet) with varying architecture (25-layer vs deeper 32/16-layer), varying Lambda (0 vs 0.05), PCA (True or False), Augmentation of the training set (True or False), Window size (F = 20 or F = 60), and training-test split ratios (70/30% vs 90/10%). To ensure that the results were not biased by specific data splits, we validated each parameter across five unique seeds for split of the data (Monte-Carlo cross validation). For each training epoch a 5-fold cross validation was also utilized. Models were evaluated on the held-out test set using accuracy & ROC-AUC to measure classification success and threshold sensitivity.

The different models were then compared using ANOVA. It was found that 25-layer architecture (p = 1.2e-07), PCA False (p = 1.4e-17), and a 90/10% split ratio (p = 1.1e-04) yielded better test accuracies. The impact of Lambda was also significant, 0.05 yielding better results in cases of larger windows (p=5.7e-04). The effect of Augmentation was not significant on the test accuracies (p=0.92) however, it did lead to better training accuracies; to avoid overtraining, augmentation was not performed for the future models.

Utilizing the parameters obtained above, multiple models were again trained with varying window length (F), ranging from 5 frames to 100 frames. It was found that the test accuracies increased until 50 frames but then saturated. The trained models were then fed in a full session, to identify the trial and non-trial sections.

Furthermore, to test that the models are learning a biological phenomenon and not some patterns in the data, 500 models (100 for each rat), were trained with shuffled labels and the best parameters. (**Supplementary Fig. 5**)

#### Statistics used

Median and 25-75% confidence intervals were utilized for reporting all the values.

To account for the repeated-measures nature of the data (multiple sessions per animal), behavioural metrics and percentage of activated/suppressed neurons were analysed using Linear Mixed-Effects (LME) models. Experimental Phase (e.g., Self-Administration, Extinction, Reinstatement) was treated as a fixed effect, while Animal Identity was included as a random intercept to account for individual variability and non-independence of observations. Time-based metrics (ITI and durations) were log-transformed prior to analysis to ensure normality of residuals. Pairwise post-hoc comparisons between phases were performed using contrast matrices within the LME framework, with significance set at α=0.05.

### Graphical User Interface based Analysis software

All the analysis can be redone, with the help of GUI script available with the data. Using these GUI scripts one can plot various graphs, for different types of cell. Explore individual cell activity over different sessions, compare it with baseline, measure p-values, and explore heatmaps generated for all the trials for a particular session. One can also recreate all the figures in the paper and change the parameters for SVM, NN, and the simulation, and run them again. (**Supplementary Fig. 8**).

## Acknowledgements

The authors gratefully acknowledge the excellent technical support by Michaela Kaiser and Marion Schmitt. We acknowledge the use of Gemini (Google) for linguistic editing and structural refinement of the manuscript text. Custom MATLAB scripts were used for data analysis. Initial code structures were refactored and optimized using Gemini (Google); however, all final outputs were manually verified by the authors for mathematical accuracy and logical consistency. We acknowledge funding by the German Research Foundation (CRC 1134 functional ensembles, project B4), DFG, 402170461-TRR265 (Spanagel et al., 2024), and ERA-Net JTC programme (project ID 01EW2402-IBRAA.

## Author Contributions

A.A., I.S., T.K., and W.S. wrote the manuscript. A.A. and I.S. analysed the data. I.S. S.P., J.B-F. performed the experiments. T.K. and W.S. supervised the project.

## Data availability

All custom MATLAB scripts, object-oriented classes, and functions utilized for neural data analysis, behavioral tracking, and alignment are publicly available on GitHub at https://github.com/tkunerlab/mPFC_DualEnsemble. This repository also includes the processed behavioral metrics, alignment datasets, and statistical summaries required to replicate the findings, alongside the exact source code used to generate all manuscript figures. Additionally, the repository features interactive data explorers to facilitate the dynamic visualization and inspection of the underlying datasets.

---

**Supplementary Fig. 1.**
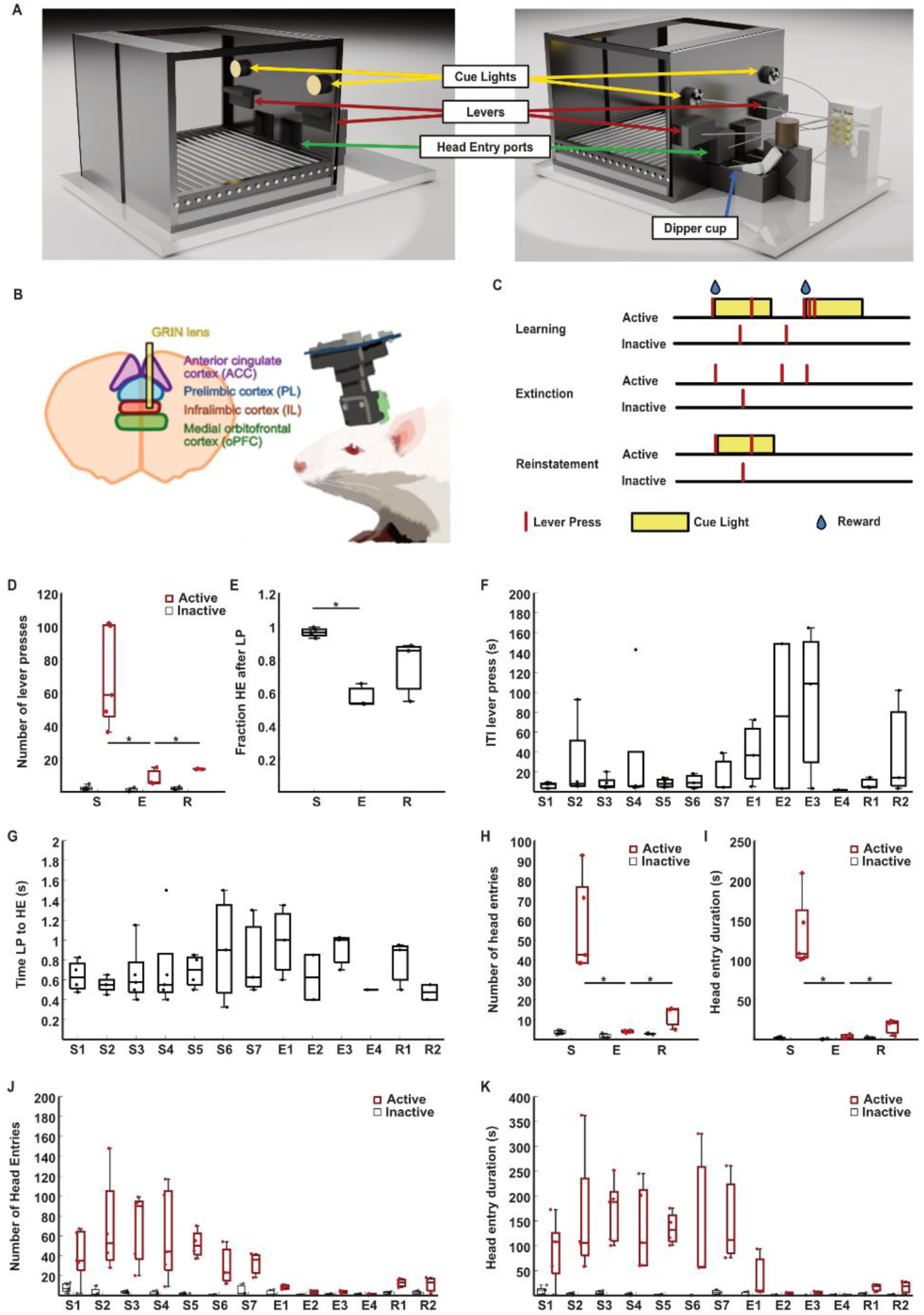
**A.** A 3d rendered model of the conditioning setup. **B.** A schematic of the mPFC, along with a sketch of rat implanted with a miniscope. **C.** Schematic of the trial structure. Yellow boxes denote cue light, red bars denote lever presses, blue drop denotes reward. **D.** Number of lever presses by the rats during different session types. **E.** Conditional probability of a head entry (HE) occurring after a lever press (LP). **F.** Intertrial interval between subsequent lever presses for all sessions. **G.** Time taken by the rat from lever press to head entry. **H-K.** Number (H,J) and duration (I,K) of head entries per session for all session types (H,I) and sessions (J,K) respectively. For the box plots, the central mark indicates the median, and the bottom and top edges of the box indicate the 25th and 75th percentiles, respectively. The whiskers extend to the most extreme data points not considered outliers. S- Self-administration, E-Extinction, R-Reinstatement. Red boxes denote the active lever press, and black boxes denote the inactive. For S, n=5; for E and R, n=3; *p<0.05; Liner-mixed effects model.

**Supplementary Figure 2.**
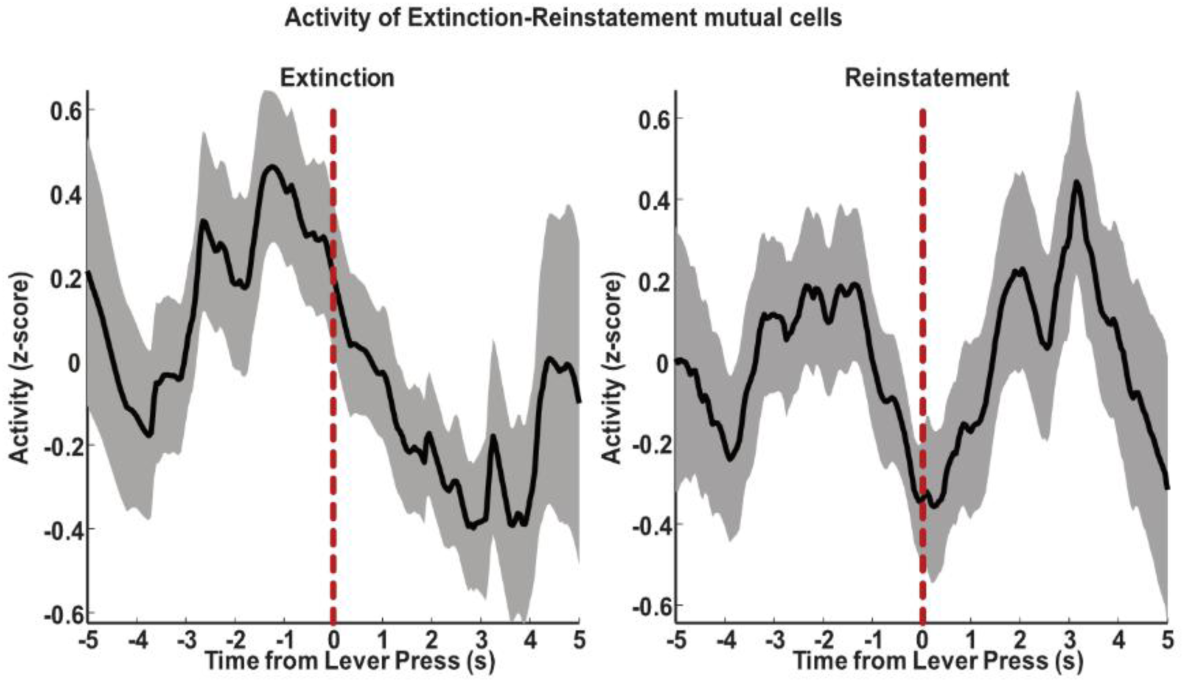
Mutually exclusive Extinction and Reinstatement neurons. Activity of the mutually exclusive neurons of Extinction and Reinstatement (n=16). Black line indicates the mean, shaded area represents SEM, red dashed line denotes lever press.

**Supplementary Figure 3:**
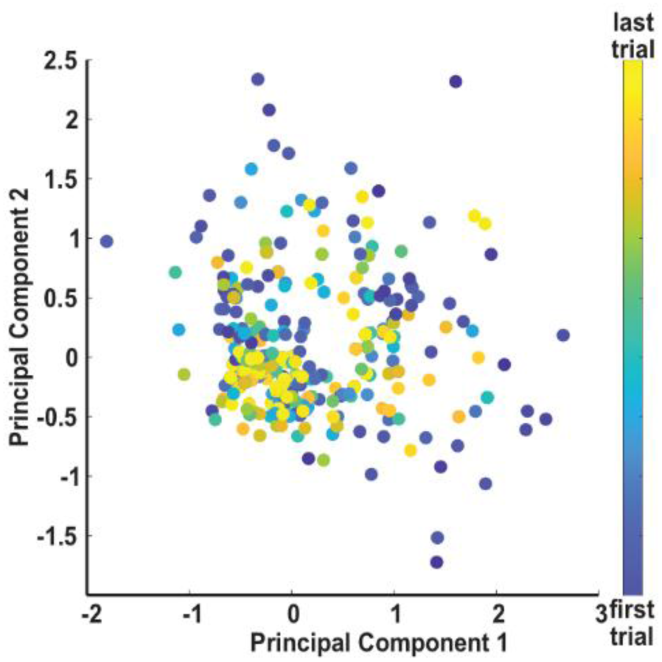
PCA for trial progression. Principal component analysis (PCA) showing the population trajectory during the first five self-administration (S) sessions. Data are projected onto the first two PCs, with trial progression represented by a colour gradient from dark blue (start/first trial) to bright yellow (end/last trial).

**Supplementary Figure 4:**
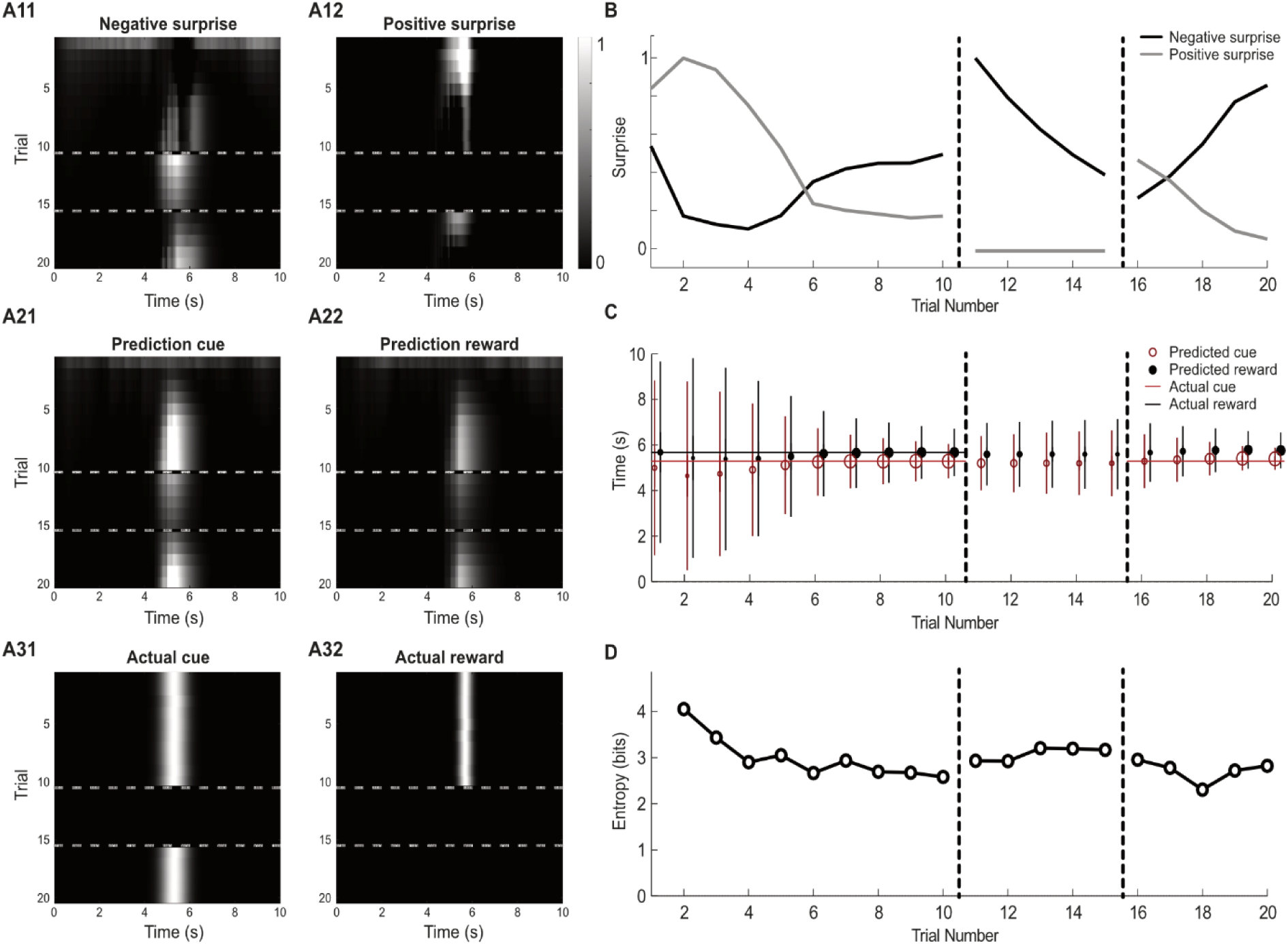
Temporal dynamics of prediction and surprise. **A:** Heatmap representations of signal intensity over time. Each row of heatmaps illustrates the temporal evolution of specific signals across successive trials (y-axis: trials; x-axis: simulated time). Colour intensity corresponds to signal magnitude, ranging from 0 (no signal, dark) to 1 (maximal activity, bright). A11–A12: Negative and positive surprise signals, A21–A22: Predicted cue and reward signals, A31–A32: Actual cue and reward signals. **B:** Comparative magnitude of surprise over trials. The line graph tracks the peak intensity of surprise signals as learning progresses. The black line denotes negative surprise, while the grey line represents positive surprise. **C:** Temporal evolution of cue (red) and reward (black) predictions. The relationship between predicted and actual events is shown across trials (x-axis: trial number; y-axis: time). The horizontal line indicates the timing of the actual cue and reward. Dots represent the predicted cue, error bars represent standard deviation of the prediction, dot size reflects prediction strength. **D:** The panel displays the calculated Shannon’s entropy for both E1 and E2 ensembles over trials.

**Supplementary Figure 5:**
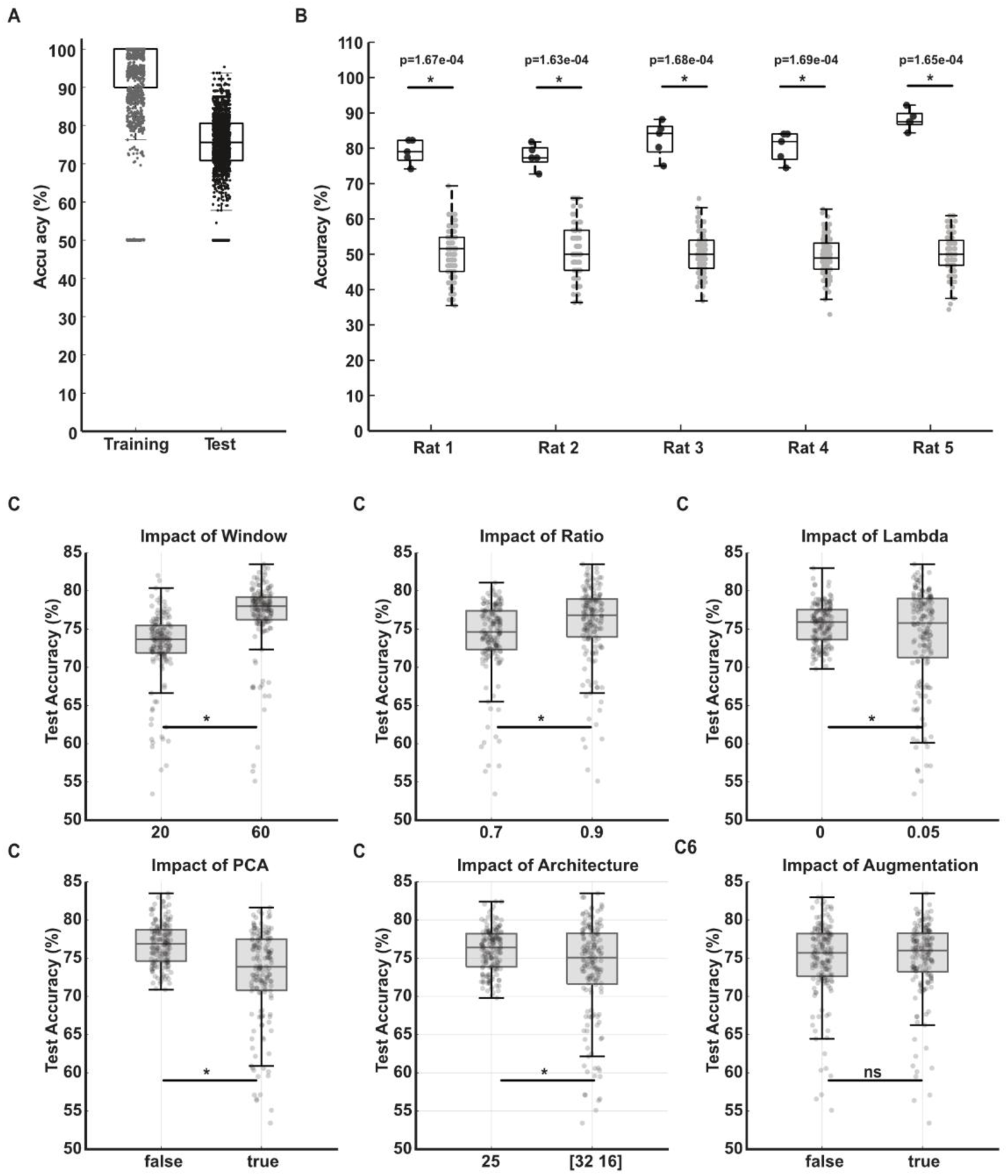
Neural Network model performances. **A:** Training and test accuracies for 4000 models trained across the whole hyperparametric space. **B:** Test accuracies for the models trained with the best-performing parameters for each rat with real labels (black) vs shuffled labels (grey). Real data: 5 models per rat, Shuffled data: 100 models per rat. **C:** Test accuracies of models trained in (A), split to understand the effect of different parameters on performance. *p<0-05

**Supplementary Figure 6.**
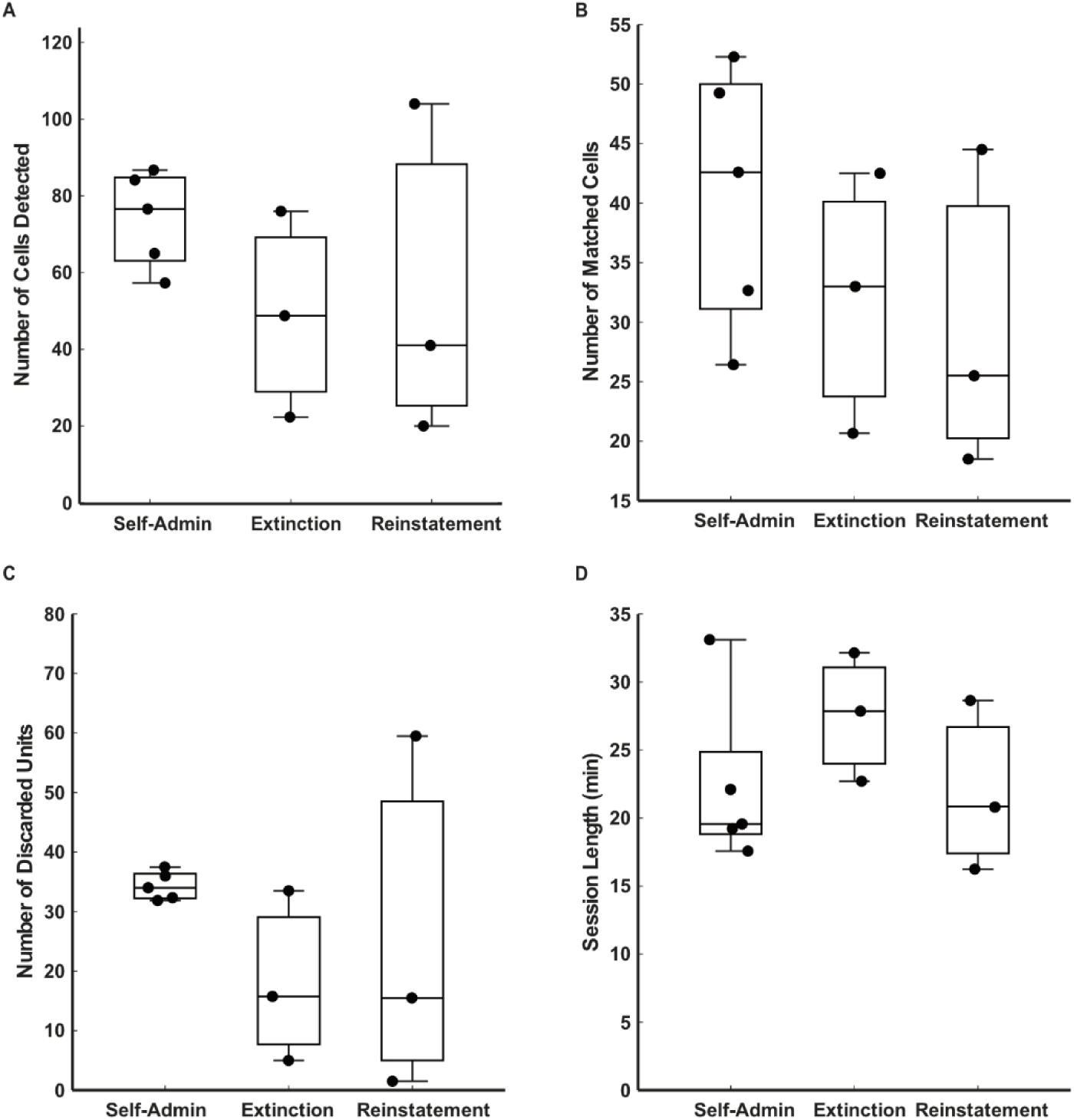
Additional information about the data. **A.** Number of cells detected per session type. **B.** Number of cells that matched per session type. **C.** Number of units that were discarded per session type. **D.** Length of the sessions recorded. The central mark indicates the median, and the bottom and top edges of the box indicate the 25th and 75th percentiles, respectively. The whiskers extend to the most extreme data points not considered outliers. Each individual scatter point represents an animal.

**Supplementary Figure 7:**
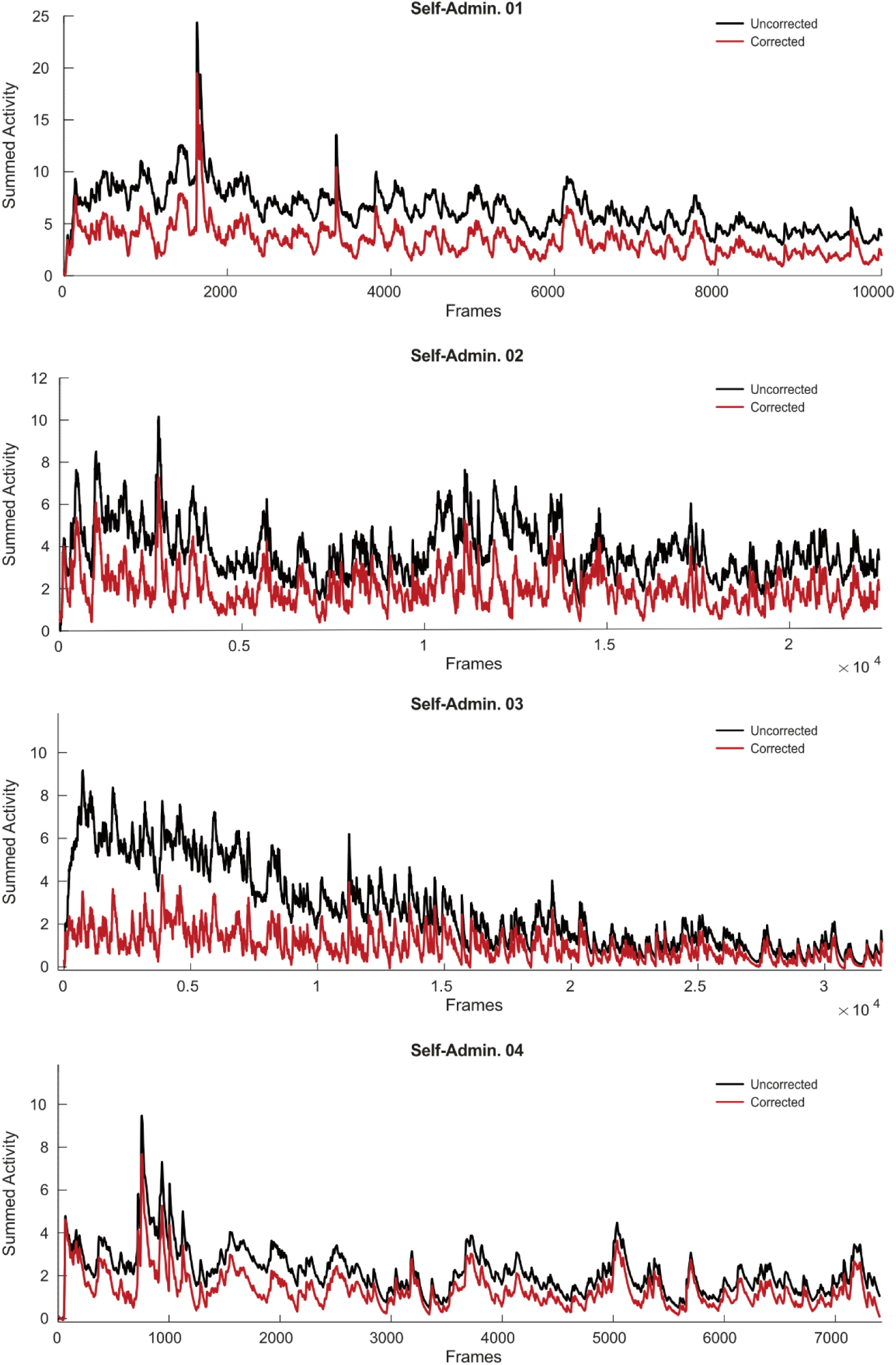
Florescence decay correction. Uncorrected and corrected traces from Rat 1 for the first four self-administration sessions. Black lines indicate uncorrected traces, whereas red lines indicate corrected traces.

**Supplementary Figure 8:**
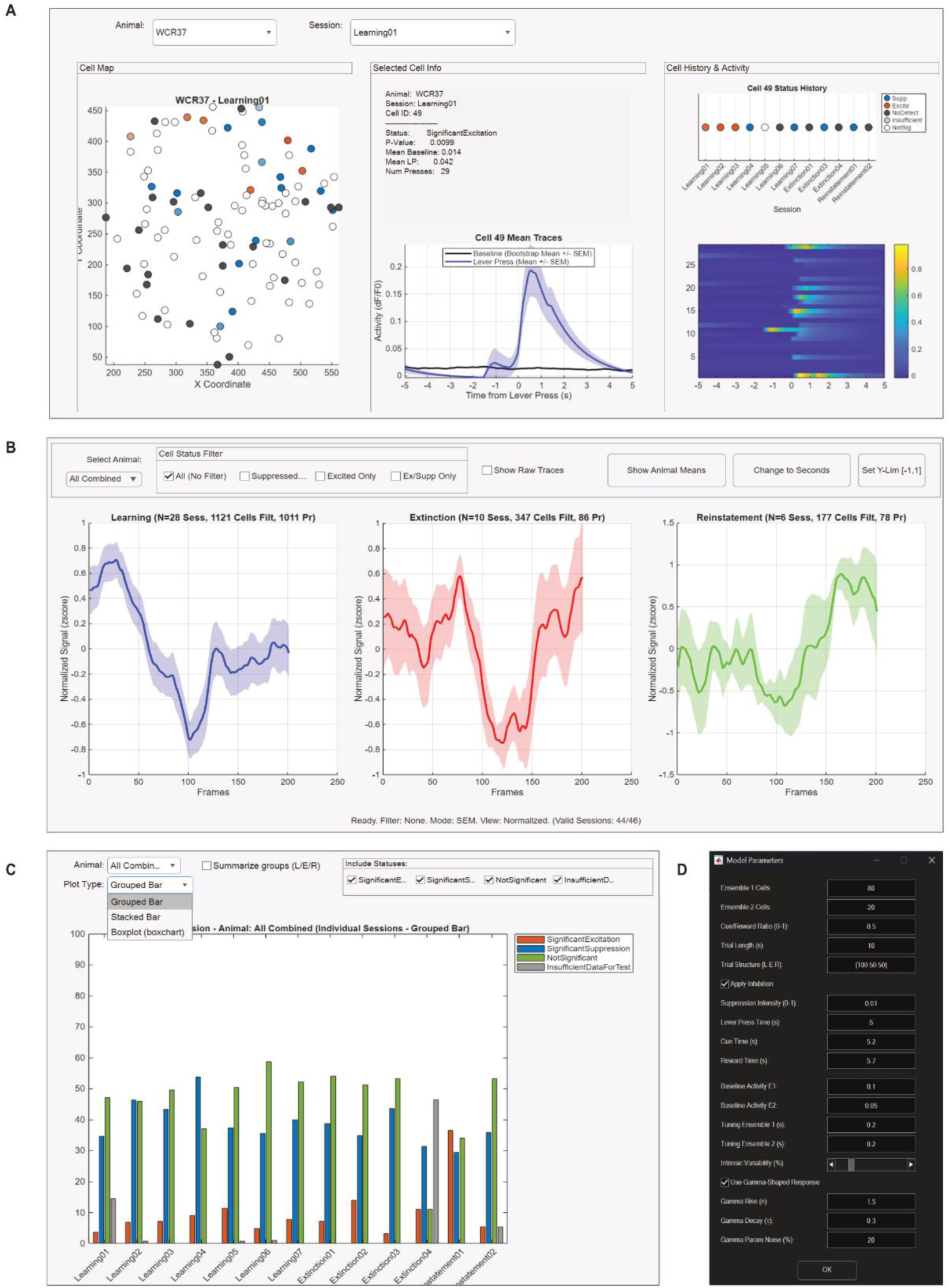
UI based data explorers and simulation. **A:** Cell activity explorer GUI. **B:** Population dynamics explorer GUI. **C:** Status Explorer GUI. **D:** Simulation parameters GUI.

## Notes

### Competing Interest Statement

The authors have declared no competing interest.

