## Supplementary figures and images for "A Dual-Ensemble Model of Infralimbic Cortex Function in Behavioural Flexibility"

### Supplementary Figure 1

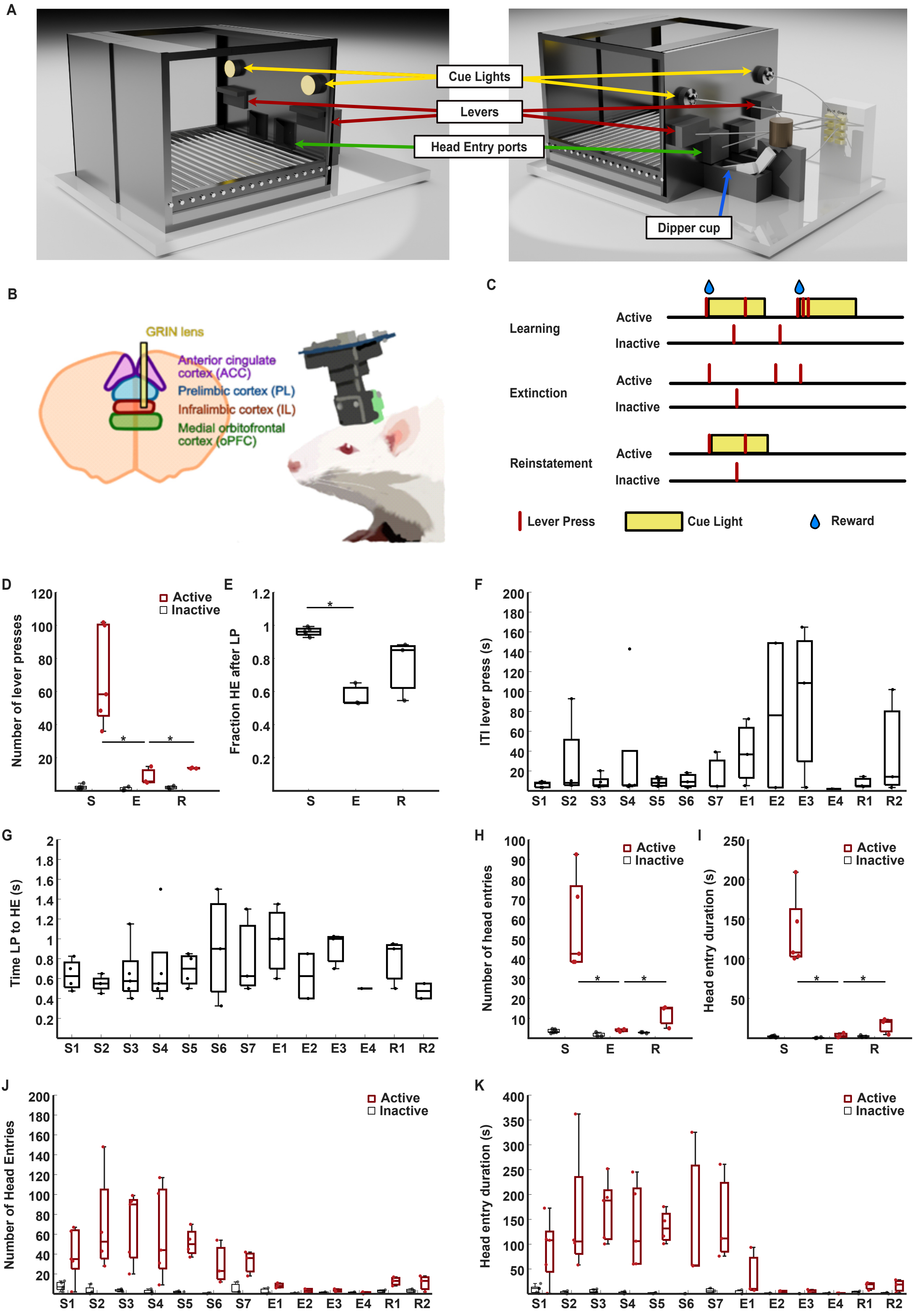

### Supplementary Figure 2

Activity of Extinction-Reinstatement mutual cells

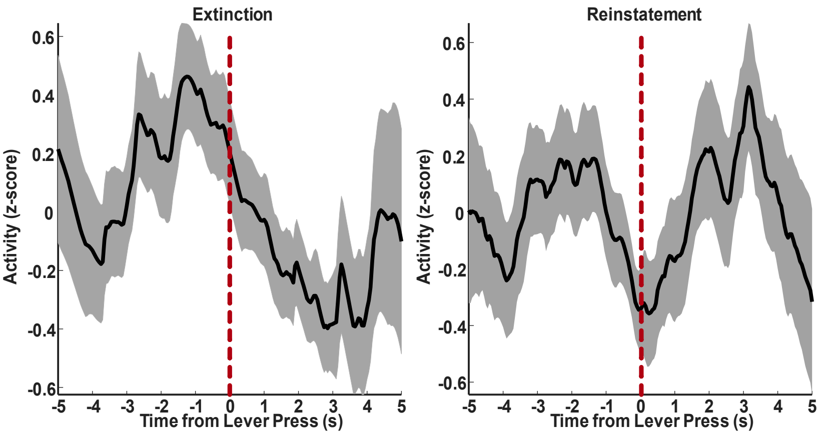

### Supplementary Figure 3

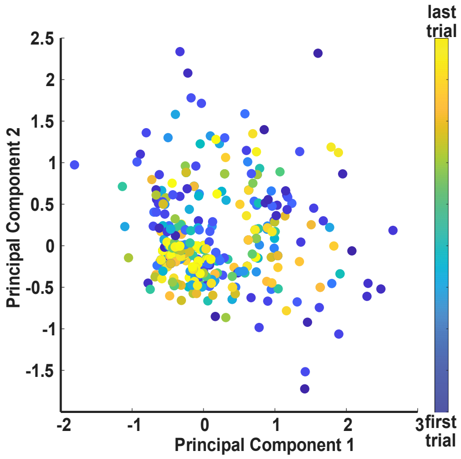

### Supplementary Figure 4

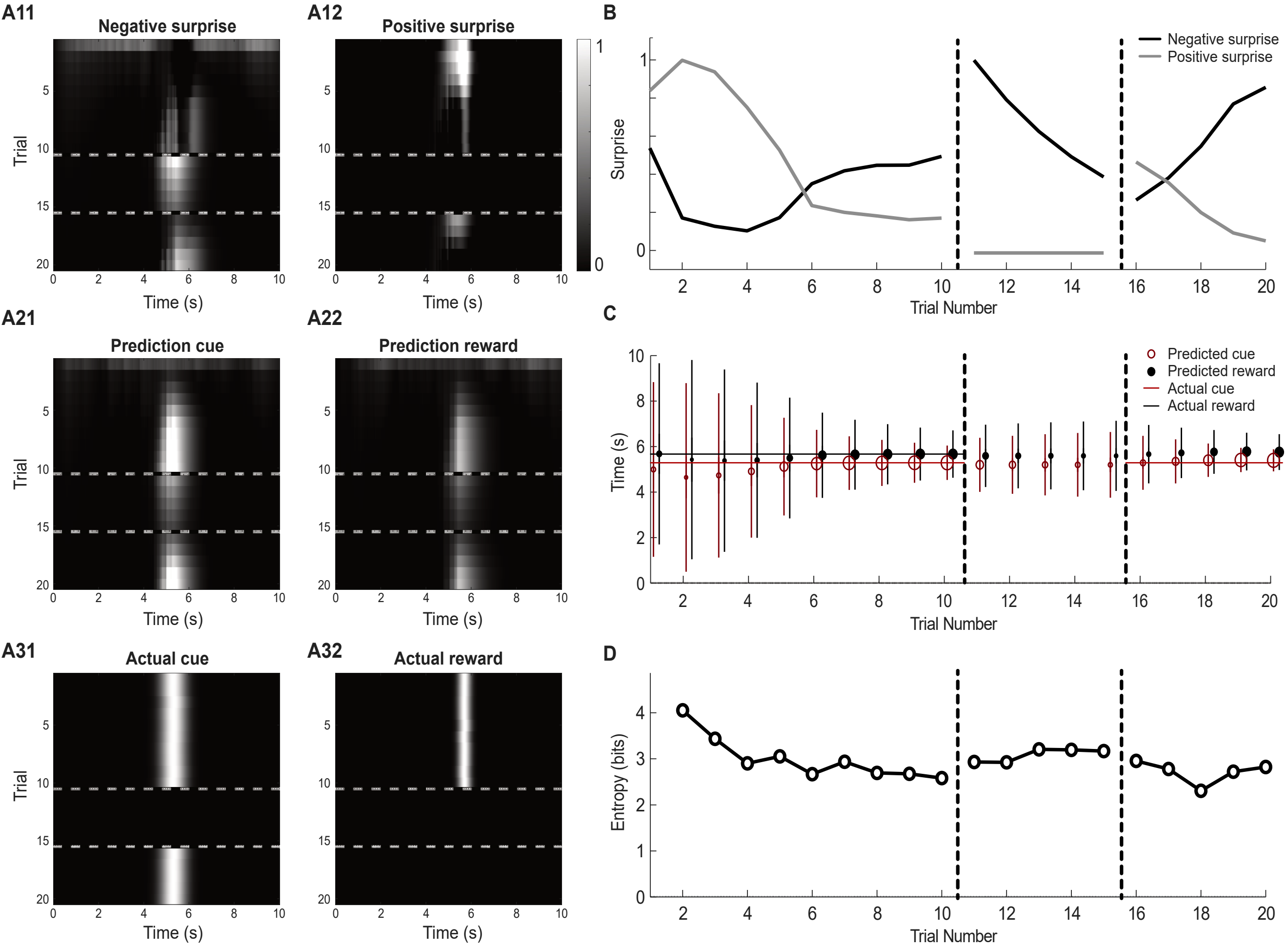

### Supplementary Figure 5

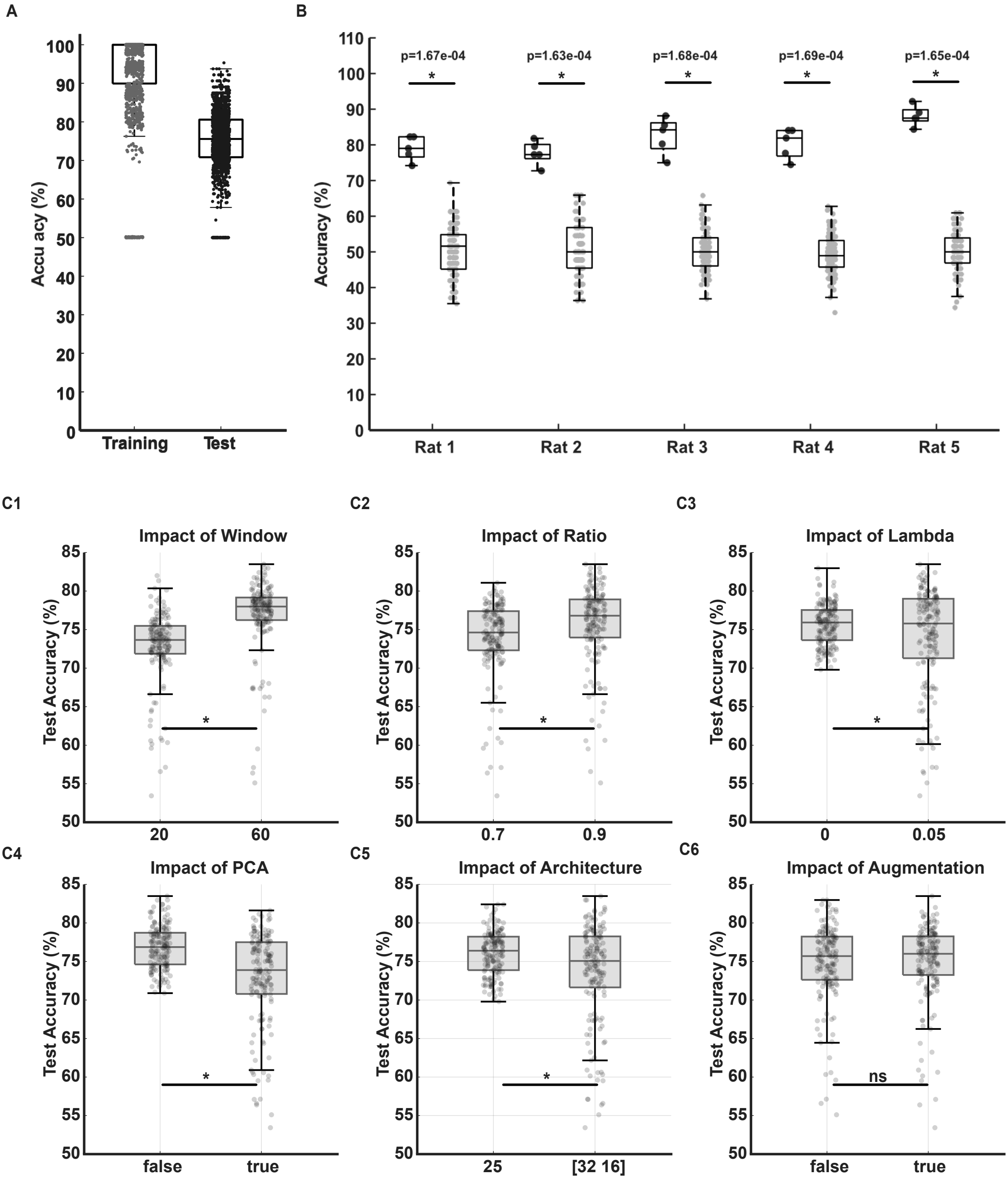

### Supplementary Figure 6

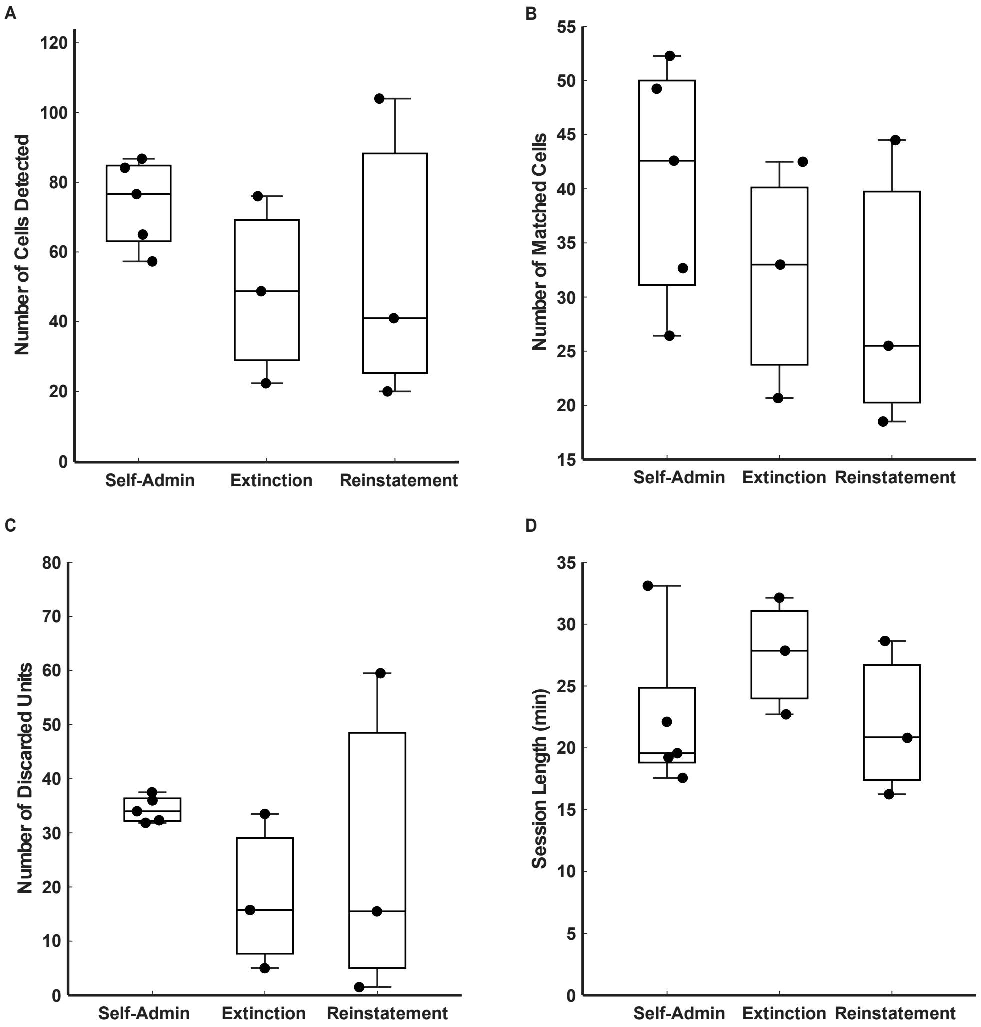

### Supplementary Figure 7

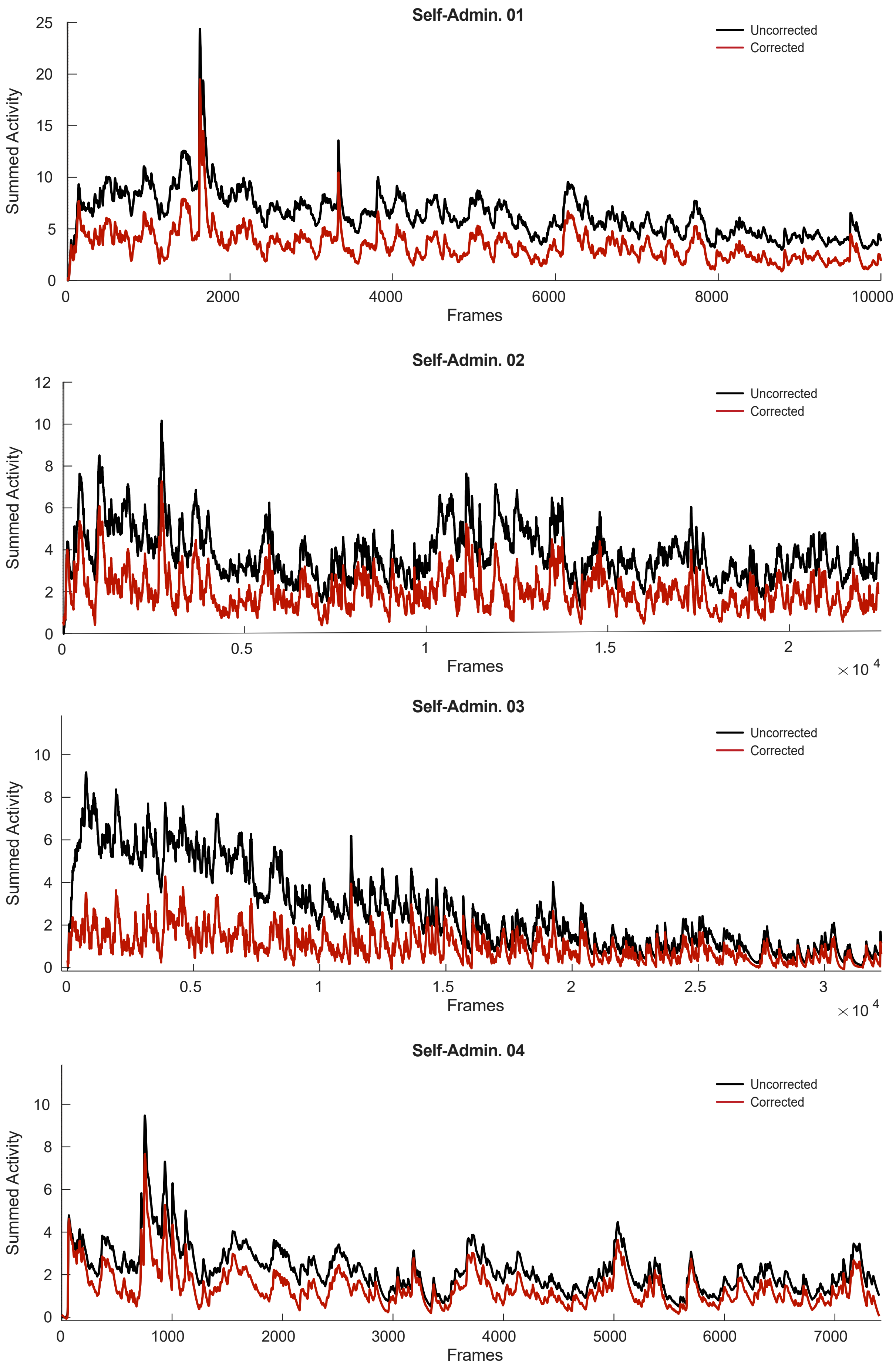

### Supplementary Figure 8

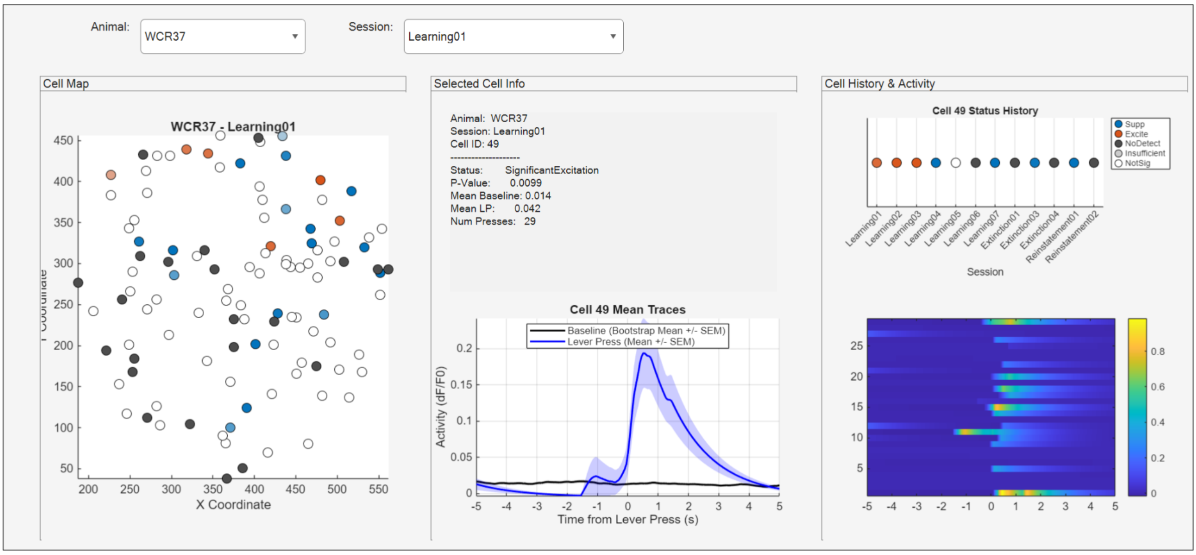

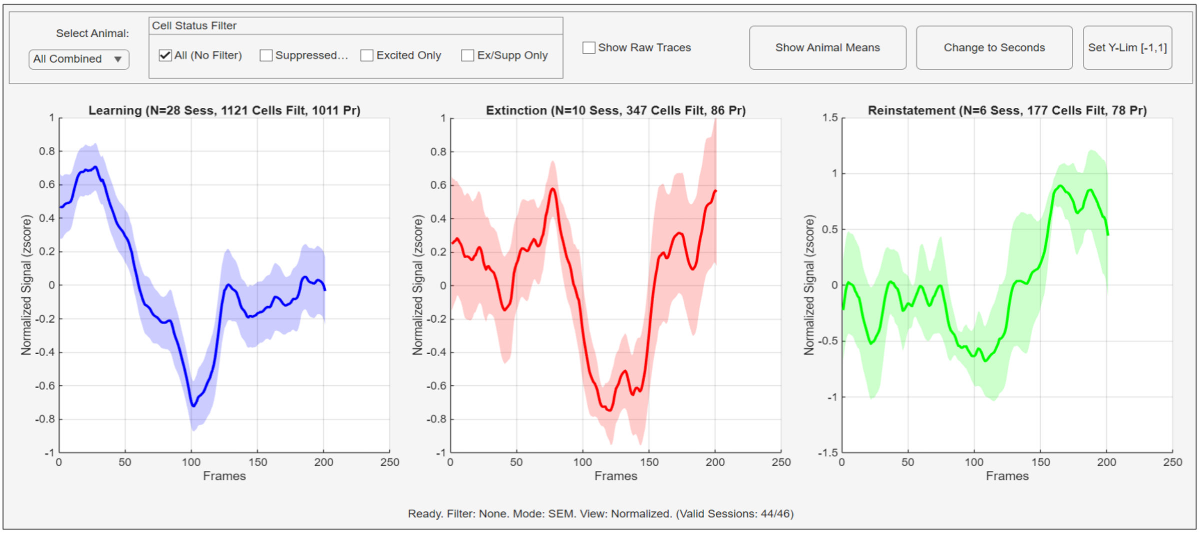

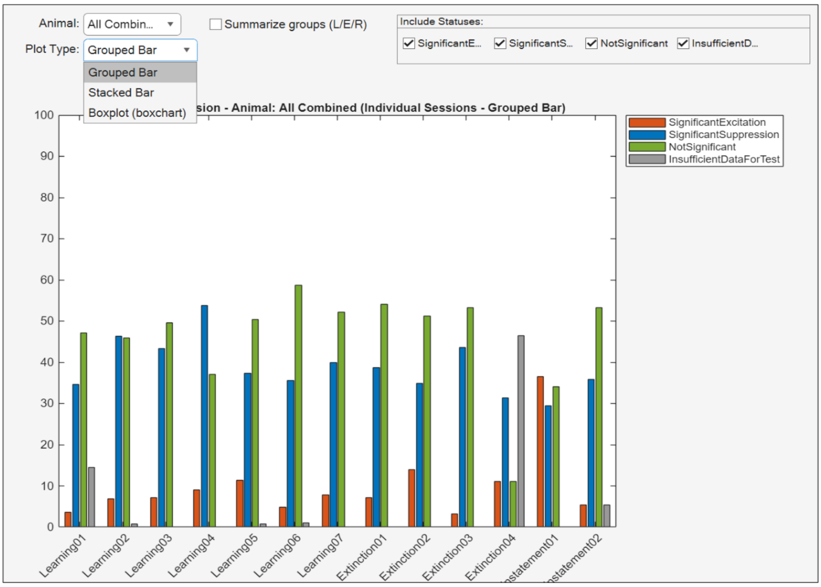

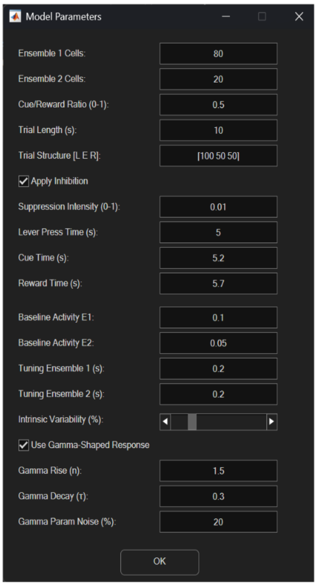
